# A common human *MLKL* polymorphism confers resistance to negative regulation by phosphorylation

**DOI:** 10.1101/2022.09.08.507056

**Authors:** Sarah E. Garnish, Katherine R. Martin, Maria Kauppi, Victoria Jackson, Rebecca Ambrose, Vik Ven Eng, Shene Chiou, Yanxiang Meng, Daniel Frank, Emma C. Tovey Crutchfield, Komal M. Patel, Annette V. Jacobsen, Georgia K. Atkin-Smith, Ladina Di Rago, Marcel Doerflinger, Christopher R. Horne, Cathrine Hall, Samuel N. Young, Vicki Athanasopoulos, Carola G. Vinuesa, Kate E. Lawlor, Ian P. Wicks, Gregor Ebert, Ashley P. Ng, Charlotte A. Slade, Jaclyn S. Pearson, Andre L. Samson, John Silke, James M. Murphy, Joanne M. Hildebrand

**Affiliations:** The Walter and Eliza Hall Institute, Parkville, VIC, 3052, Australia; University of Melbourne, Department of Medical Biology, Parkville, 3052, Australia; Centre for Innate Immunity and Infectious Diseases, Hudson Institute of Medical Research, Clayton, VIC, 3168, Australia; Department of Molecular and Translational Science, Monash University, Clayton, VIC, 3168, Australia; Department of Microbiology, Monash University, Clayton, 3168, VIC, Australia; University of Melbourne, Faculty of Medicine, Dentistry and Health Sciences, Parkville, 3052, Australia; Department of Immunology and Infection, John Curtin School of Medical Research, Australian National University, ACT, Australia; Institute of Virology, Technical University of Munich/Helmholtz Munich, Munich, Germany; Clinical Haematology Department, The Royal Melbourne Hospital and Peter MacCallum Cancer Centre, Parkville, 3052, Australia; Department of Clinical Immunology & Allergy, Royal Melbourne Hospital, Parkville, VIC, 3052, Australia

## Abstract

Across the globe, 2-3% of humans carry the *p.Ser132Pro* single nucleotide polymorphism in *MLKL*, the terminal effector protein of the inflammatory form of programmed cell death, necroptosis. We show that this substitution confers a gain in necroptotic function in human cells, with more rapid accumulation of activated MLKL^S132P^ in biological membranes and MLKL^S132P^ overriding pharmacological and endogenous inhibition of MLKL. In mouse cells, the equivalent *Mlkl S131P* mutation confers a gene dosage dependent reduction in sensitivity to TNF-induced necroptosis in both hematopoietic and non-hematopoietic cells, but enhanced sensitivity to IFN-β induced death in non-hematopoietic cells. *In vivo*, *Mlkl^S131P^* homozygosity reduces the capacity to clear *Salmonella* from major organs and retards recovery of hematopoietic stem cells. Thus, by dysregulating necroptosis, the S131P substitution impairs the return to homeostasis after systemic challenge. Present day carriers of the *MLKL S132P* polymorphism may be the key to understanding how MLKL and necroptosis modulate the progression of complex polygenic human disease.

## INTRODUCTION

Necroptosis is a caspase independent form of programmed cell death that originated as a defense against pathogens ^1, 2, 3, 4, 5^. Highly inflammatory in nature, necroptosis results in the permeabilization of biological membranes and the release of cytokines, nucleic acids, and intracellular proteins into the extracellular space ^6^. Necroptosis is induced by danger- or pathogen-associated molecular patterns that signal via transmembrane receptors or intracellular pattern recognition receptors ^7, 8, 9, 10, 11^. Of the various initiating stimuli, the most well studied necroptotic pathway is downstream of tumor necrosis factor receptor 1 (TNFR1)^12^. In physiological contexts that favor low cellular inhibitor of apoptosis protein 1 (cIAP1) and caspase-8 activity, TNFR1 signals culminate in the formation of a high molecular weight platform called the necrosome that is nucleated by heterooligomeric RIPK1 and RIPK3 ^13, 14, 15^. Here, the terminal executioner protein, MLKL, is phosphorylated and activated by its upstream kinase, RIPK3 ^16, 17, 18^. Following phosphorylation, MLKL dissociates from RIPK3, oligomerizes, and is trafficked to biological membranes where it interacts with Phosphatidylinositol Phosphates (or ‘PIPs’) ^19, 20, 21, 22, 23, 24, 25, 26, 27, 28, 29^. In human cells, association of activated MLKL oligomers with biological membranes can be inhibited by the synthetic compound necrosulfonamide ^16, 22, 28^ or inhibitory phosphorylation of MLKL at Serine 83 ^30^. Pharmacological or mutation driven disruption at any major necroptotic signaling checkpoint compromises a cell’s capacity to execute necroptosis.

In mouse studies, MLKL-mediated cell death has been implicated as a driver or suppressor of diseases spanning almost all physiological systems depending on the pathological context. The generation of *Mlkl* gene knockout (*Mlkl^-/-^*), knock-in and conditional knockout mouse models have enabled the role of necroptosis in infectious and non-infectious challenges to be dissected in physiological detail ^31^. Interestingly, genetic deletion of *Mlkl* has no overt developmental or homeostatic effects, with the exception of a reduction in age-related sterile inflammation in female mice ^18, 32, 33^. This is in direct contrast with two mouse models harboring *Mlkl* point mutations that dysregulate MLKL activation, *Mlkl^D139V^* and *Mlkl^S83G^*, which exhibit early neonatal death and severe inflammatory phenotypes ^30, 34^. Altogether, these observations suggest that while constitutive absence of MLKL-mediated death is benign, imbalanced execution of necroptotic cell death is deleterious.

Consistent with this notion, more than 20 unique disease-associated human germline gene variants in the core necroptotic machinery, encompassing *RIPK1, RIPK3, MLKL*, have been identified ^35, 36^. In one family, a haplotype including a rare *MLKL* loss-of-function gene variant (*p.Asp369GlufsTer22*, rs561839347) is associated with a severe and progressive novel neurogenerative spectrum disorder characterized by global brain atrophy ^37^. A more frequent *MLKL* loss-of-function gene variant (*p.Gln48Ter*, rs763812068) was found to be >20 fold enriched in a cohort of Hong Kong Chinese patients suffering from Alzheimer’s disease ^38^ and common variants that cluster around the MLKL brace region were shown to be enriched *in trans* in a cohort of chronic recurrent multifocal osteomyelitis patients ^34^. More recently, a hypomorphic *MLKL* missense gene variant (*p.G316D*, rs375490660) was reported to be associated with Maturity Onset Diabetes of the Young ^39^.

Here we present the cellular and physiological characterization of a serine to proline missense polymorphism at MLKL amino acid 132 (*p.Ser132Pro*; *S132P*). The *S132P* polymorphism is the third most frequent human *MLKL* missense coding variant in the gnomAD database; a large repository of whole genome and exome sequence data from humans of diverse ancestry ^40^. To examine the potential human disease-causing effects of this *MLKL* variant, we exogenously expressed *MLKL^S132P^* in human cell lines and introduced the mouse counterpart variant (*Mlkl^S131P^*) into a genetically modified mouse model, revealing that this polymorphism confers MLKL gain-of-function in a cell- and stimulus-dependent manner. This MLKL gain-of-function manifests in *in vivo* changes to the immune response, impaired bacterial clearance, and defective emergency hematopoiesis. These observed phenotypes provide important insights into how this highly frequent human *MLKL S132P* polymorphism may contribute to the progression of complex disease.

## RESULTS

### Carriers of the *p.Ser132Pro* polymorphism exhibit diverse inflammatory disease profiles

With a global minor allele frequency (MAF) of 0.0138, the *MLKL S132P* (rs35589326) polymorphism is predicted to be carried by 2-3% of the human population. It has not been detected in individuals assigned East Asian ancestry, is rare in individuals of African or Latino/Admixed American ancestry and is carried by an estimated 6-7% of individuals of Ashkenazi Jewish ancestry (MAF 0.0315) (www.gnomAD.com, February 2023) **(Figure 1A)**. Notably, Ser132 is highly conserved across species **(Figure 1B)**.

**Figure 1.**
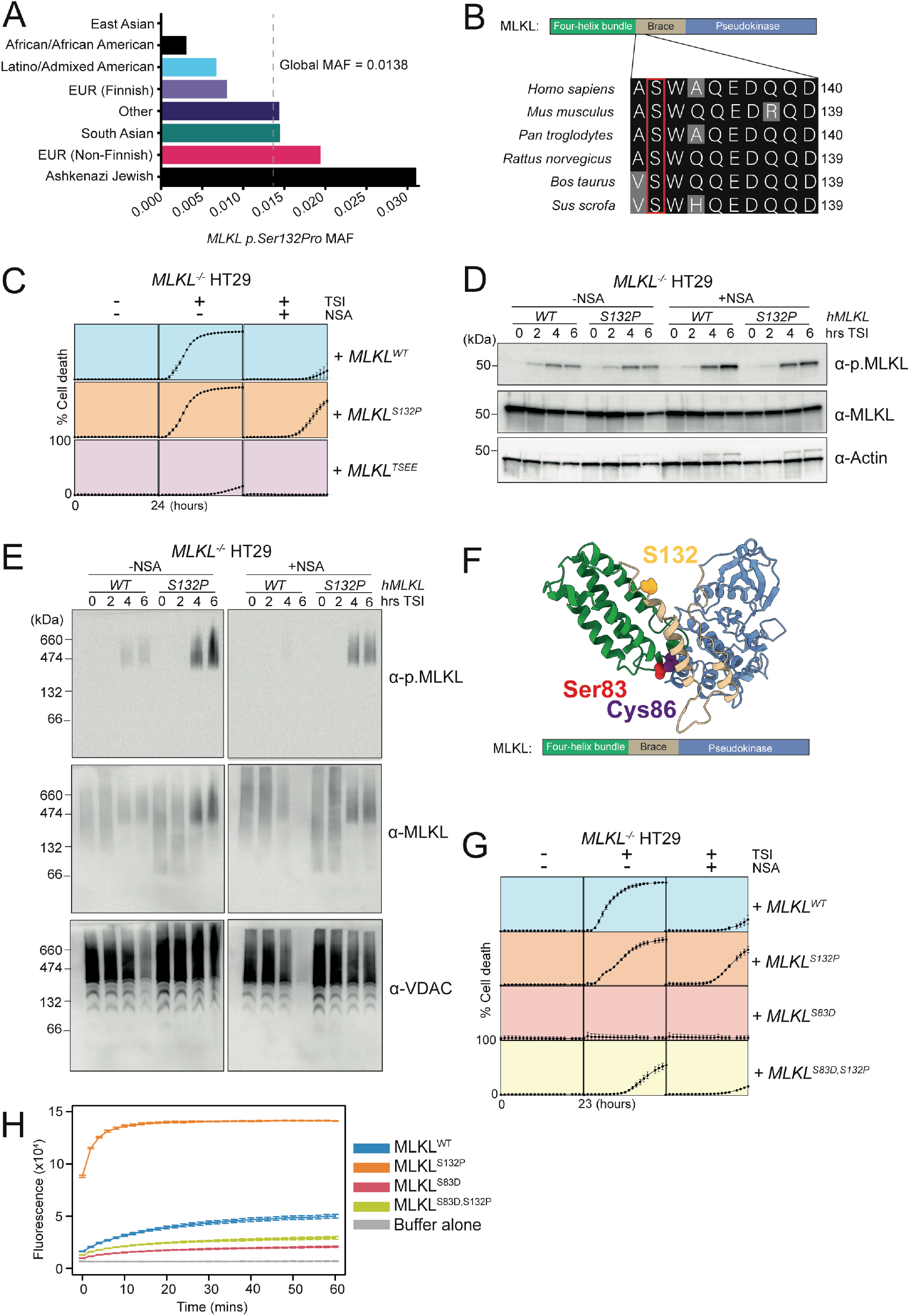
MLKL^S132P^ executes cell death in the presence of necrosulfonamide or Ser83 inhibition. **(A)** Minor allele frequency of *MLKL p.Ser132Pro* according to the gnomAD database, stratified by ancestry. **(B)** Sequence alignment of conserved serine in the MLKL orthologs across different species. **(C)** MLKL^WT^, MLKL^S132P^ and MLKL^TSEE^ expression was induced in *MLKL^-/-^* HT29 cells with 100 ng/ml doxycycline (Dox) and treated with necroptotic stimulus (TNF, Smac mimetic, IDN-6556; TSI) in the presence or absence of MLKL inhibitor Necrosulfonamide (NSA; 1µM). Cell death was measured every hour for 24 hours by percentage of SYTOX Green positive cells quantified using IncuCyte SX5 live cell imaging. Independent cell lines were assayed in *n=3-9* experiments, with errors bars indicating the mean **±** SEM. **(D)** Western blot analyses of whole cell lysates taken 3h post TSI stimulation in the presence or absence of NSA from doxycycline (100 ng/ml) induced *MLKL^-/-^* HT29 cells expressing *MLKL^WT^* or *MLKL^S132P^*. **(E)** High molecular weight phosphorylated MLKL^S132P^ is present at enhanced levels in crude membrane fractions following Blue-Native PAGE under TSI stimulation in the presence or absence of NSA. **(F)** S83 (red), C86 (purple) and S132 (gold) highlighted as spheres on cartoon representation of human MLKL. Four-helix bundle domain shown in green, brace helices shown in beige and pseudokinase domain shown in blue. Homology model is generated from PDB:2MSV and PDB: 4MWI of human MLKL, which were aligned using the full-length murine crystal structure (PDB:4BTF)^18, 74, 75^. **(G)** *MLKL S132P* mutation reconstitutes necroptotic signaling in the presence of S83 inhibitory phosphorylation (MLKL^S83D^). Human MLKL expression was induced in *MLKL^-/-^* HT29 cells with doxycycline (100 ng/ml) and treated TSI in the presence or absence of NSA (1µM). Cell death was measured every hour for 23 hours by percentage of SYTOX Green positive cells quantified using IncuCyte SX5 live cell imaging. Independent cell lines were assayed in *n=4* experiments, with errors bars indicating the mean **±** SEM. **(H)** Liposome dye release assays using 0.5 µM recombinant full-length MLKL^WT^, MLKL^S132P^, MLKL^S83D^ and MLKL^S83D,S132P^. Release of 5(6)-Carboxyfluorescein was measured by fluoresence (485 nm excitation wavelength, 535 nm emission wavlength) every 2 min over 60 min. Data represent mean **±** SD of triplicate measurements, representative of three independent assays. **(D/E)** Blot images are representative of at least three independent experiments.

Two heterozygous carriers of the *MLKL S132P* polymorphism were identified in an Australian registry of patients suffering from immune related disease. Patient 1, a female of South American heritage, was diagnosed with SAPHO (synovitis, acne, pustulosis, hyperostosis, osteitis) syndrome (inheritance chart unavailable). Patient 2, a female of European heritage, was diagnosed with systemic IgG4 disease **(Supplementary Figure 1A)**. Both patients have one or more immediate family members carrying the *MLKL S132P* polymorphism who were also diagnosed with inflammatory diseases in early adulthood. Following whole genome sequencing, Patient 1 was found to exhibit a region of loss of heterozygosity (5:96031569 – 5:96364063) which covers *CAST, ERAP1, ERAP2* and *LNPEP.* These genes have been previously associated with inflammatory disease ^41, 42, 43, 44, 45^. Patient 2 does not carry any other predicted pathogenic gene variants. Primary peripheral blood mononuclear cells (PBMCs) isolated from patient 2 showed reduced MLKL protein levels, accompanied by a nominal increase in pro-inflammatory cytokine production in response to the Toll-like receptor (TLR) agonists, lipopolysaccharide and poly I:C, relative to age and sex matched healthy donor PBMCs not carrying the *MLKL S132P* polymorphism (**Supplementary Figure 1B, C)**.

### *MLKL^S132P^* confers resistance to chemical and natural regulatory inhibition

To examine any changes to MLKL function conferred by this common polymorphism, we stably transduced parental wild-type (WT) and *MLKL^-/-^* versions of the human colonic epithelial cell line HT29 with doxycycline inducible *MLKL^WT^*and *MLKL ^S132P^* gene expression constructs. We also included the T357/S358 phosphosite mutant *MLKL^TSEE^*, previously shown to be inactive^20, 46^, as a negative control for necroptosis induction. Cells expressing *MLKL^S132P^* died with similar kinetics to *MLKL^WT^* following necroptotic stimulation (TNF, Smac mimetic compound A and pan-caspase inhibitor IDN-665; TSI) and, as expected, *MLKL^TSEE^* reconstituted cells were resistant to necroptotic cell death **(Figure 1C, Supplementary Figure 1D, E)**. Interestingly, in the presence of the MLKL inhibitor necrosulfonamide (NSA), TSI-stimulated cells expressing *MLKL^S132P^* exhibited higher levels of cell death than their *MLKL^WT^*counterparts at later timepoints **(Figure 1C, Supplementary Figure 1D)**. We termed this override of MLKL inhibition ‘breakthrough’ cell death. Breakthrough death also occurred when *MLKL^S132P^* was co-expressed with endogenous *MLKL* and increased when *MLKL^S132P^* expression was augmented by higher doses of doxycycline **(Figure 1C**, **Supplementary Figure 1D, E).** We also observed breakthrough death in both wild-type and *MLKL^-/-^* forms of the human monocytic cell line U937, indicating this is not a cell type specific phenomenon **(Supplementary Figure 1F, G)**. Breakthrough death was neither due to differences in MLKL expression, nor changes in MLKL phosphorylation (T357/S358), because both were equivalent between *MLKL^WT^* and *MLKL^S132P^* reconstituted cells **(Figure 1D, Supplementary Figure 1H, I)**. Instead, cellular fractionation experiments suggest that breakthrough death was likely due to enhanced association of MLKL^S132P^ with biological membranes **(Figure 1E, Supplementary Figure 1J)**.

The small molecule inhibitor, necrosulfonamide (NSA), blocks human MLKL activity through covalent modification of Cysteine 86, located in the MLKL four helix-bundle, executioner domain **(Figure 1F)** ^16, 47^. Recently, it was discovered that MLKL can be endogenously phosphorylated at the proximal residue, Serine 83, and this phosphorylation event plays a species conserved inhibitory role in both human (S83) and mouse (S82) **(Figure 1F)** ^30^. Like NSA, phosphorylation of MLKL S83 does not alter the capacity of RIPK3 to phosphorylate MLKL at S357/T358 but prevents necroptosis by blocking the association of MLKL with cellular membranes ^30^. To test whether phosphorylation at S83 is, like NSA, less effective at inhibiting MLKL^S132P^ induced death, we created stable HT29 (*MLKL^-/-^* and wild-type) cell lines that exogenously express gene constructs encoding *MLKL^WT^*, *MLKL^S132P^*, *MLKL^S83D^* (phosphomimetic) and double mutant *MLKL^S83D,S132P^*. Consistent with recent studies^30^, the phosphomimetic mutation of S83, *MLKL^S83D^*, ablated the capacity of MLKL to execute necroptotic cell death **(Figure 1G)**. Also, as previously reported, the inhibitory effect of *MLKL^S83D^* overrode the activating effects of endogenous MLKL phosphorylation at Ser357/Thr358 **(Figure 1G, Supplementary Figure 1K, L)**. Exogenous expression of *MLKL^S83D^* also reduced total levels of necroptotic cell death in the presence of endogenous MLKL, indicating that this S83 phosphomimetic acts in a dominant negative manner **(Supplementary Figure 1L)**. Strikingly, combining the MLKL^S132P^ and MLKL^S83D^ substitutions to create the compound mutant MLKL^S83D, S132P^ restored necroptotic cell death responsiveness, albeit with reduced kinetics and reduced maximal cell death when compared to MLKL^S132P^ alone **(Figure 1G)**. The *MLKL^S132P^* variant also partially overcomes the dominant negative effect of S83 phosphomimetic mutation in cells that endogenously express wild-type *MLKL* **(Supplementary Figure 1L)**.

To further dissect how the MLKL^S132P^ substitution facilitates gain-of-function, we used liposome dye release assays to test the membrane damaging capacity of recombinant full-length MLKL^WT^, MLKL^S132P^, MLKL^S83D^ and MLKL^S83D, S132P^ expressed and purified from insect cells. We found that the membrane damaging capacity of recombinant MLKL^S132P^ was increased, and MLKL^S83D^ reduced, relative to MLKL^WT^ **(Figure 1H)**. Consistent with our observations in cells, the combination mutant MLKL^S83D, S132P^ displayed a membrane damaging capacity greater than MLKL^S83D^ alone but reduced in comparison to MLKL^WT^ **(Figure 1H)**. Our results suggest S132P promotes membrane association, and this is likely to contribute to the gain-of-function we observed *in vitro* under pharmacological and natural inhibition.

Given the clear gain-of-function conferred to MLKL in the context of simulated inhibitory phosphorylation (MLKL^S83D, S132P^), we questioned whether mutations that resemble this phosphomimetic mutant may occur naturally. According to the gnomAD database (www.gnomAD.com, Jan 2023), no individuals have been recorded with a polymorphism that encodes the *p.Ser83Asp* (S83D) replacement. However, there are individuals that carry closely related changes, *p.Ser83Cys* (MAF 3.54e10^-5^) and *p.Arg82Ser* (MAF 1.57e10^-5^). We created stable *MLKL^-/-^* HT29 cell lines that exogenously express gene constructs encoding *MLKL^S83C^* and *MLKL^R82S^* and assessed their capacity to execute necroptosis. Cells expressing *MLKL^S83C^* died with similar kinetics to *MLKL^WT^* following necroptotic stimulation however, *MLKL^R82S^* reconstituted cells were resistant to cell death **(Supplementary Figure 1M)**. Consistent with our observations for *MLKL^S83D^*, *MLKL^R82S^* was phosphorylated at Ser357/Thr358, and the *MLKL^R82S,S132P^* compound mutant restored necroptotic killing function **(Supplementary Figure 1M, N).** Whilst gene variants at or adjacent to the S83 inhibitory phosphorylation site are less frequent than the *MLKL S132P* polymorphism in humans, they nonetheless strengthen the precedent for genetically encoded diversity in human MLKL function, and the potential for functionally synergistic or neutralising combinations of *MLKL* gene variants.

### *MLKL^S131P^* mice exhibit differences in steady state immune cell populations

To address if the *MLKL S132P* polymorphism contributes to immunoinflammatory disease, we performed detailed histological, immunophenotypic, and experimental analyses of a CRISPR generated knockin mouse which carried the orthologous mutation, *Mlkl^S131P^*. *Mlkl^S131P^* heterozygotes and homozygotes were born according to the expected Mendelian ratios and had normal lifespans **(Figure 2A, B)**. The healthy presentation of *Mlkl^S131P^* mice contrasts with the lethal phenotypes of mice engineered to encode *Mlkl^D139V^* and *Mlkl^S83G^* mutations ^30, 34^. Wild-type, *Mlkl^S131P^* heterozygote and homozygote mice had comparable body weight and no gross histological differences up to 9 months of age **(Figure 2C, Supplementary Figure 2A)**. Notably, distinct from *Mlkl^D139V^* or *Mlkl^S83G^* homozygotes, no inflammation was detected in the salivary glands, mediastinum, liver or lungs (**Supplementary Figure 2A)** ^30, 34^. No differences in the number of circulating platelets, red blood cells, or white blood cells were observed between wild-type, *Mlkl^S131P^* heterozygote and homozygote mice across age **(Supplementary Figure 2B-D)**. However, reduced numbers of classical Ly6C^hi^ ‘inflammatory’ monocytes in the bone marrow, but not in the secondary lymphoid organs (spleen and inguinal lymph nodes), were observed in *Mlkl^S131P^* homozygotes and heterozygotes relative to wild-type littermate controls (**Figure 2D, Supplementary Figure 2E, J, O)**. In the secondary lymphoid organs, *Mlkl^S131P^* homozygotes had a small but significant increase splenic CD4^+^ T cells and B cells relative to wild-type littermate controls **(Figure 2E, Supplementary Figure 2M, N).** Compared to wild-type controls, *Mlkl^S131P^* heterozygotes had significant increases in splenic and lymph node B cell populations, as well as splenic CD8^+^ T cells and Ly6C^lo^ monocytes **(Figure 2E, F, Supplementary Figure 2J-S)**. All other innate and adaptive immune cell populations were comparable between genotypes in the bone marrow, spleen, and inguinal lymph nodes **(Figure 2D-F, Supplementary Figure 2E-S)**. Further, wild-type, heterozygote and homozygote mice had comparable levels of plasma cytokines at steady state **(Figure 2G)**.

**Figure 2.**
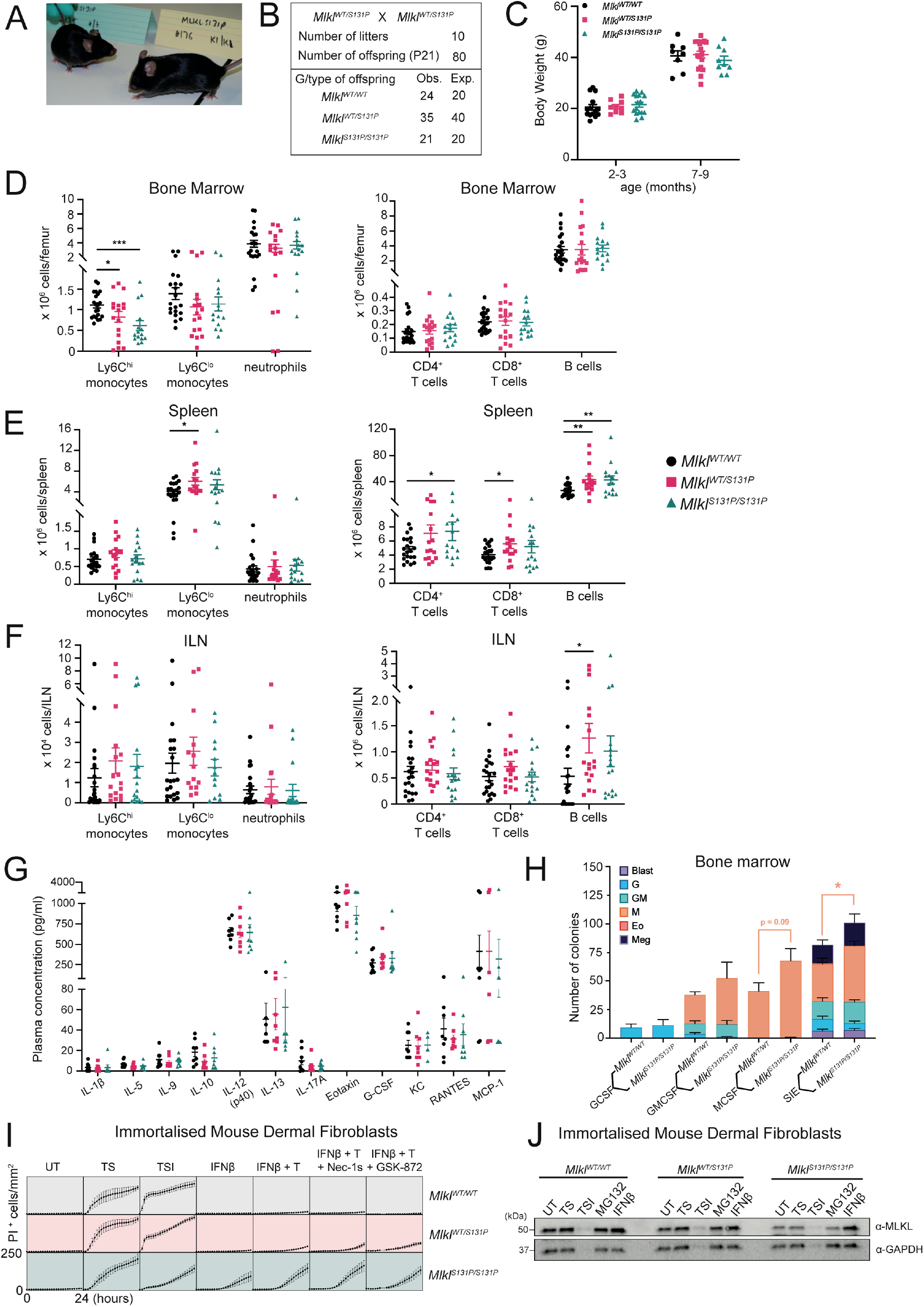
In mice, *Mlkl^S131P^* homozygosity is tolerated but results in steady state immune cell population differences. **(A)** Macroscopic appearance of *Mlkl^WT/WT^* and *Mlkl^S131P/S131P^* mice at 8-12 weeks of age. **(B)** *Mlkl^S131P/S131P^* mice are born according to Hardy-Weinberg equilibrium as observed in the distribution of genotypes from *Mlkl^WT/S131P^* heterozygous intercrosses. **(C)** Body weight of *Mlkl^WT/WT^*, *Mlkl^WT/S131P^* and *Mlkl^S131P/S131P^* mice at 3-4 or 7-9 months of age. Each dot represents an individual mouse (*n= 8-17,* mean ± SEM). **(D-F)** Flow cytometry quantification of innate (Ly6C^hi^ monocytes, Ly6C^lo^ monocytes and neutrophils) and adaptive (CD4^+^ T cells, CD8^+^ T cells and B cells) in the bone marrow **(D)**, spleen **I** and inguinal lymph nodes **(F)** of 8–12-week-old *Mlkl^WT/WT^, Mlkl^WT/S131P^,* and *Mlkl^S131P/S131P^* mice. Each symbol represents one individual mouse sampled and error bars represent mean ± SEM for *n=7-11* mice as indicated. **(G)** Multiplex measurement of plasma cytokines from 6-12 week old mice. Each symbol represents one individual mouse sampled, with mean ± SEM of *n=*4-8. **(H)** Type and number of colonies from 25,000 unfractionated bone marrow cells cultured in G-CSF (10^3^ U/mL), GM-CSF (10^3^ U/mL), SIE [SCF (100 ng/mL), IL-3 (10 ng/mL), EPO (2 U/mL)] were scored after 7 days. Error bars represent mean ± SEM for *n* =4 mice per genotype. **(I, J)** Immortalized mouse dermal fibroblasts (MDFs) were isolated from *Mlkl^WT/WT^, Mlkl^WT/S131P^,* and *Mlkl^S131P/S131P^* mice and stimulated as indicated for 24 hours for quantification of PI-positive cells using IncuCyte S3 live cell imaging **(I)** or for 6 hours for western blot analysis **(J)**. Death data represent mean ± SEM for independently generated cell lines of *n*=3-5. *p<0.05, **p<0.01 calculated using an unpaired, two-tailed Students t-test.

To investigate the underlying cause of reduced Ly6C^hi^ monocytes in the bone marrow of *Mlkl^S131P^* homozygote mice, we examined both the abundance of different myeloid stem cell populations, and their capacity to differentiate and form colonies *ex vivo*. *Mlkl^WT/WT^* and *Mlkl^S131P/S131P^* hematopoietic stem cells exhibited equivalent capacity to generate blast, eosinophil, granulocyte, granulocyte-macrophage, and megakaryocyte colonies under all stimulations investigated **(Figure 2H)**. While the number and composition of myeloid progenitor cell populations in the bone marrow were not significantly different **(Supplementary Figure 2T, U)**, compared to *Mlkl^WT/WT^*, *Mlkl^S131P/S131P^* bone marrow gave rise to an increased number of macrophage colonies under SCF, IL-3 and EPO combined stimulation **(Figure 2H)**.

We next sought to address whether mouse MLKL^S131P^ exhibited a gain-in-function comparable to our finding for MLKL^S132P^ in human cells. Primary and immortalized fibroblasts isolated from the dermis of *Mlkl^S131P^* homozygote, heterozygote and wild-type mice were examined for their relative capacity to die in the presence or absence of death stimuli. We did not observe any differences in TNF-induced apoptosis (TNF and Smac mimetic; TS) between *Mlkl^S131P/S131P^* and *Mlkl^WT/WT^* immortalized MDFs **(Figure 2I)**. However, *Mlkl^S131P/S131P^* MDFs showed a diminished capacity to undergo TSI-induced (TNF, Smac mimetic compound A, pan-caspase inhibitor IDN-6556; TSI) necroptotic cell death **(Figure 2I, Supplementary Figure 2V)**. This was driven by reduced MLKL protein levels in MDF cells with the *Mlkl^S131P^* allele **(Figure 2J, Supplementary Figure 2W)**. Immortalized MDFs, and to a lesser extent primary MDFs, did show clear *Mlkl^S131P^* allele-dependent sensitivity to IFN-β, a strong inducer of *Mlkl* gene expression in mice **(Figure 2I, Supplementary Figure 2V)** ^48^. This sensitivity was further enhanced by TNF and was refractory to inhibitors of RIPK1 (Nec-1s) or RIPK3 (GSK′872) **(Figure 2I, Supplementary Figure 2V)**. These results are analogous to cells expressing the constitutively-active *MLKL^D139V^* mutant ^34^ and, together, support the notion that MLKL^S131P^ has constitutive RIPK3-independent activity when endogenously expressed in mouse cells ^34^. These endogenous *Mlkl^S131P^* observations were not limited to fibroblasts. In bone marrow derived macrophages (BMDMs), *Mlkl^S131P/S131P^* cells also exhibited diminished sensitivity to TSI-induced necroptosis and endogenously produced MLKL^S131P^ was present at reduced levels in comparison to MLKL^WT^ **(Supplementary Figure 2X, Y)**. In contrast to immortalized dermal fibroblasts, BMDMs derived from *Mlkl^S131P^* homozygotes did not undergo cell death in the presence of IFN-β alone, despite clear IFN-β induced upregulation of MLKL protein relative to untreated cells **(Supplementary Figure 2X, Y)**. Overall, these data show that, across different cell types, MLKL^S131P^ is not toxic at steady state, but upon different stimuli, both gain- and loss-of-sensitivity to necroptotic cell death is evident. This suggests that any functional deficits or enhancements in carriers of the *MLKL S132P* polymorphism are likely to be diverse, with cell- and/or context-specific manifestations.

### Emergency hematopoiesis is defective in *Mlkl^S131P/S131P^* mice

Differences in steady state immune cell populations suggest that overt phenotypes may develop in *Mlkl^S131P^* homozygotes following experimental challenge. Specifically, reduced numbers of steady state Ly6C^hi^ monocytes in the bone marrow of *Mlkl^S131P^*homozygotes could indicate a defect in hematopoiesis. To test this, we examined recovery from myelosuppressive irradiation as an assessment of hematopoietic function in mice carrying the *Mlkl^S131P^* allele. Following myelosuppressive irradiation, recovery of hematopoietic cell numbers and circulating peripheral blood cells was significantly delayed in *Mlkl ^S131P/S131P^* mice compared with wild-type controls (**Figure 3A-F)**. In *Mlkl^S131P/S131P^* mice, peripheral red blood cell and platelet numbers were significantly reduced at 14 days post irradiation, with the former also decreased at 21 days **(Figure 3A, B)**. Despite equivalent total white blood cell numbers at 21 days post irradiation, a significant reduction in peripheral monocyte numbers and a non-significant decrease trend in neutrophils was observed in *Mlkl ^S131P/S131P^*mice compared to wild-type controls **(Figure 3C, Supplementary Figure 3A, B)**. These reductions in circulating blood cell numbers were accompanied by significant decreases in the nucleated viable, progenitor, and LSK cell populations in the bone marrow of *Mlkl^S131P/S131P^* mice **(Figure 3D-F)**. Contrastingly, at 21 days post irradiation, *Mlkl ^S131P^* heterozygotes displayed a significant increase in their total nucleated viable cells in comparison to wild-type controls. This was driven predominantly by significant increases in the number of LSK cells (**Figures 3D, E)**. Impaired recovery of *Mlkl^S131P/S131P^* LSK and progenitor cell numbers was characterized by increased expression of ROS and Annexin V at 21 days post irradiation **(Supplementary Figure 3C, D)**. *Mlkl^S131P^* homozygotes had increased plasma G-CSF levels at 14 and 21 days post irradiation when compared to *Mlkl^WT/WT^*controls **(Figure 3G)**. At 21 days, all other plasma cytokines, with exception of IL-1β and RANTES, were equivalent between genotypes **(Supplementary Figure 3E)**. *Mlkl^WT/S131P^* and *Mlkl^S131P/S131P^* mice had decreased plasma IL-1β levels, whilst *Mlkl^S131P/S131P^* mice alone had reduced RANTES levels compared to *Mlkl^WT/WT^*controls **(Supplementary Figure 3E)**.

**Figure 3.**
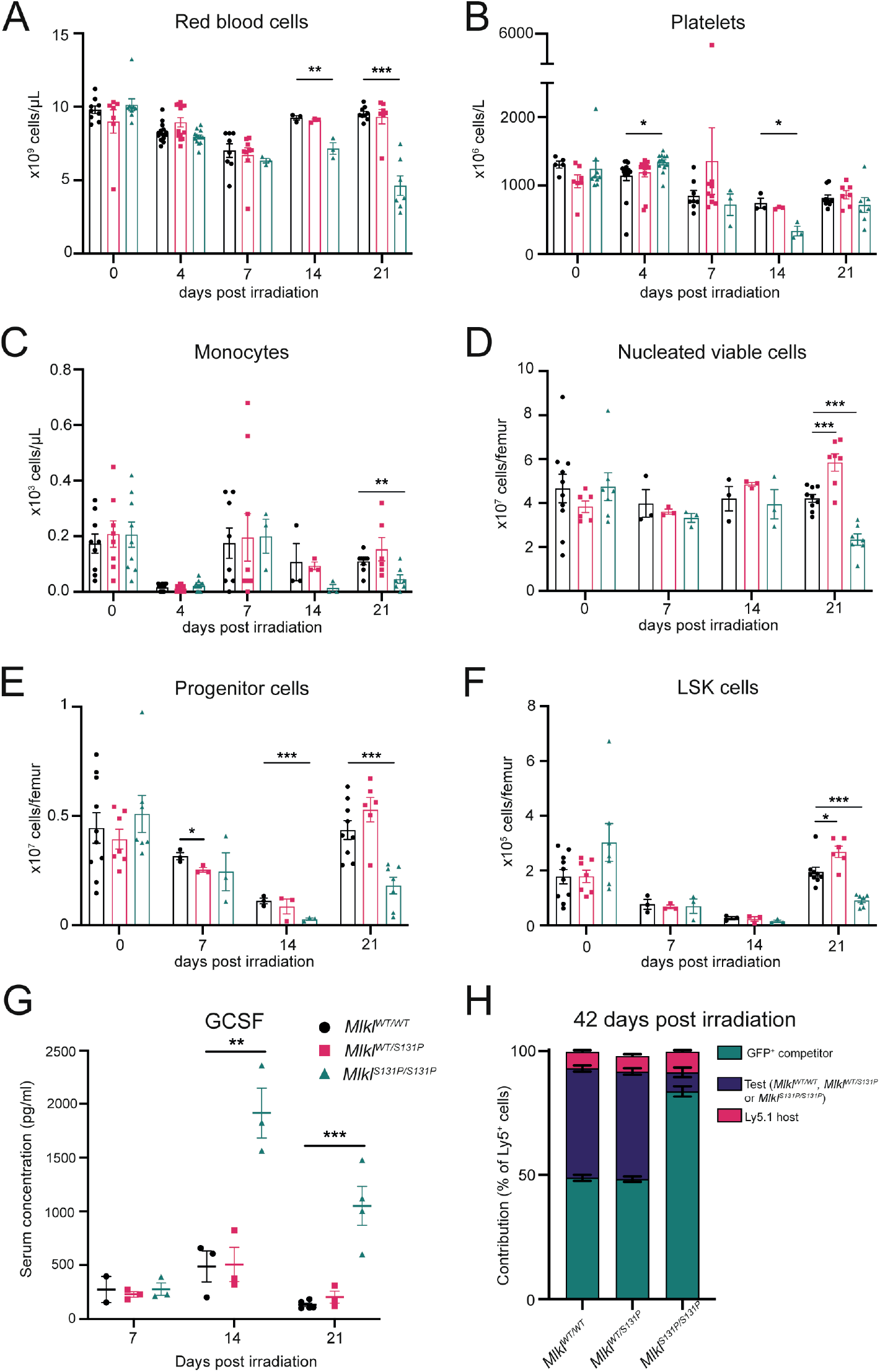
*Mlkl^S131P^* mice show delayed recovery from myelosuppressive irradiation. Peripheral red blood cells **(A)**, platelets **(B)** and monocytes **(C)** in *Mlkl^WT/WT^, Mlkl^WT/S131P^,* and *Mlkl^S131P/S131P^* mice following treatment with 5.5 Gy radiation. Quantified nucleated viable cells **(D)**, progenitor **(E)** and LSK **(F)** populations in the bone marrow of mice after myelosuppressive radiation. **(G)** Multiplex measurement of plasma G-CSF levels at 7, 14 and 21 days post-myelosuppressive radiation. Each symbol represents one individual mouse sampled, with mean ± SEM of *n=*2-5 independent mice from two separate experiments. Bone marrow from *Mlkl^WT/WT^, Mlkl^WT/S131P^* or *Mlkl^S131P/S131P^* mice on CD45^Ly5.2^ background was mixed with wild-type GFP^+^ competitor bone marrow on a CD45^Ly5.2^ background and transplanted into irradiated CD45^Ly5.1^ recipients. **(H)** Relative donor contribution to PBMCs was assessed at 6 weeks post-transplantation. Mean shown of n=5-11 recipients, with each donor bone marrow placed into 2-3 recipients. Host contribution (CD45^Ly5.1^) depicted in pink, GFP competitor in green and test (*Mlkl^WT/WT^, Mlkl^WT/S131P^* or *Mlkl^S131P/S131P^*) in purple. *p<0.05, **p<0.01, ***p<0.001 calculated using an unpaired, two-tailed Students t-test **(A-G)**.

To investigate whether the *Mlkl^S131P^* homozygote defect was intrinsic to the hematopoietic stem cells, we performed competitive transplantation studies. *Mlkl^WT/WT^*, *Mlkl^WT/S131P^* or *Mlkl^S131P/S131P^* bone marrow was mixed with GFP^+^ competitor bone marrow in a 50:50 ratio and injected into Ly5.1 irradiated hosts. Six weeks post-transplantation, *Mlkl^WT/WT^* bone marrow had competed with GFP^+^ bone marrow effectively, whilst *Mlkl^S131P/S131P^* bone marrow performed poorly, contributing to 7%, 5% and 9% of PBMCs (Ly5^+^), red blood cells and platelets respectively **(Figure 3H, Supplementary Figure 3F, G)**. Under competitive transplant conditions, *Mlkl^WT/S131P^*bone marrow competed comparably to *Mlkl^WT/WT^* with approximately 50% of the peripheral cells generated from these donor stem cells **(Figure 3H, Supplementary Figure 3F, G)**. When mixed in excess at 70:30 with GFP^+^ competitor bone marrow, *Mlkl^S131P/S131P^*stem cells were again outcompeted contributing to less than 10 % of the peripheral blood cells **(Supplementary Figure 3H-J)**. Thus, the defect in emergency hematopoiesis was intrinsic to *Mlkl^S131P/S131P^* hematopoietic stem cells, and reminiscent of an intrinsic defect previously reported for the *Mlkl^D139V^* autoactivating mutant ^34^.

### Recruitment of Ly6C^hi^ monocytes to sites of sterile inflammation is reduced in *Mlkl^S131P/S131P^* mice

To examine if the defects in emergency hematopoiesis displayed by *Mlkl^S131P/S131P^*mice could result in defective immune cell recruitment to sites of inflammation or infection, we employed a model of localized sterile inflammation induced by zymosan. Intra-peritoneal injection of zymosan results in a rapid influx of immune cells, predominantly neutrophils, into the peritoneal cavity over 72 hours ^49^. The cellular content of the peritoneal cavity and numbers of circulating peripheral blood cells was examined at 4 and 24 hours post-zymosan injection. *Mlkl^S131P/S131P^* mice displayed equivalent early recruitment of neutrophils and other immune cells to the peritoneum **(Figure 4A, B, Supplementary Figure 4A-H).** However, at 24 hours post-zymosan injection, *Mlkl^S131P/S131P^*mice had significant reductions in the number of Ly6C^hi^ monocytes recruited to the peritoneum **(Figure 4C, D, Supplementary Figure 4I-P)**. In the peripheral blood, monocytes were significantly elevated in *Mlkl^S131P/S131P^* mice in comparison to wild-type controls at 4 hours post-zymosan **(Figure 4E, Supplementary Figure 4Q-S)**. However, by 24 hours, the number of peripheral blood monocytes was equivalent between genotypes, with only a non-significant increase in neutrophils observed in *Mlkl^S131P/S131P^* mice **(Figure 4F, Supplementary Figure 4T-V)**. We also measured cytokines present in the peritoneum at 4 and 24 hours post-zymosan injection **(Figure 4G, Supplementary Figure 4W-Y)**. At 4 hours, *Mlkl^S131P/S131P^*mice had increased quantities of IL-13, IL-17A and MCP-1, whilst *Mlkl^WT/S131P^* had increased quantities of eotaxin, when compared to wild-type controls **(Figure 4G)**. No statistically significant differences were observed in the quantities of peritoneal cytokines between any genotypes at 24 hours post-injection **(Supplementary Figure 4Y)**.

**Figure 4.**
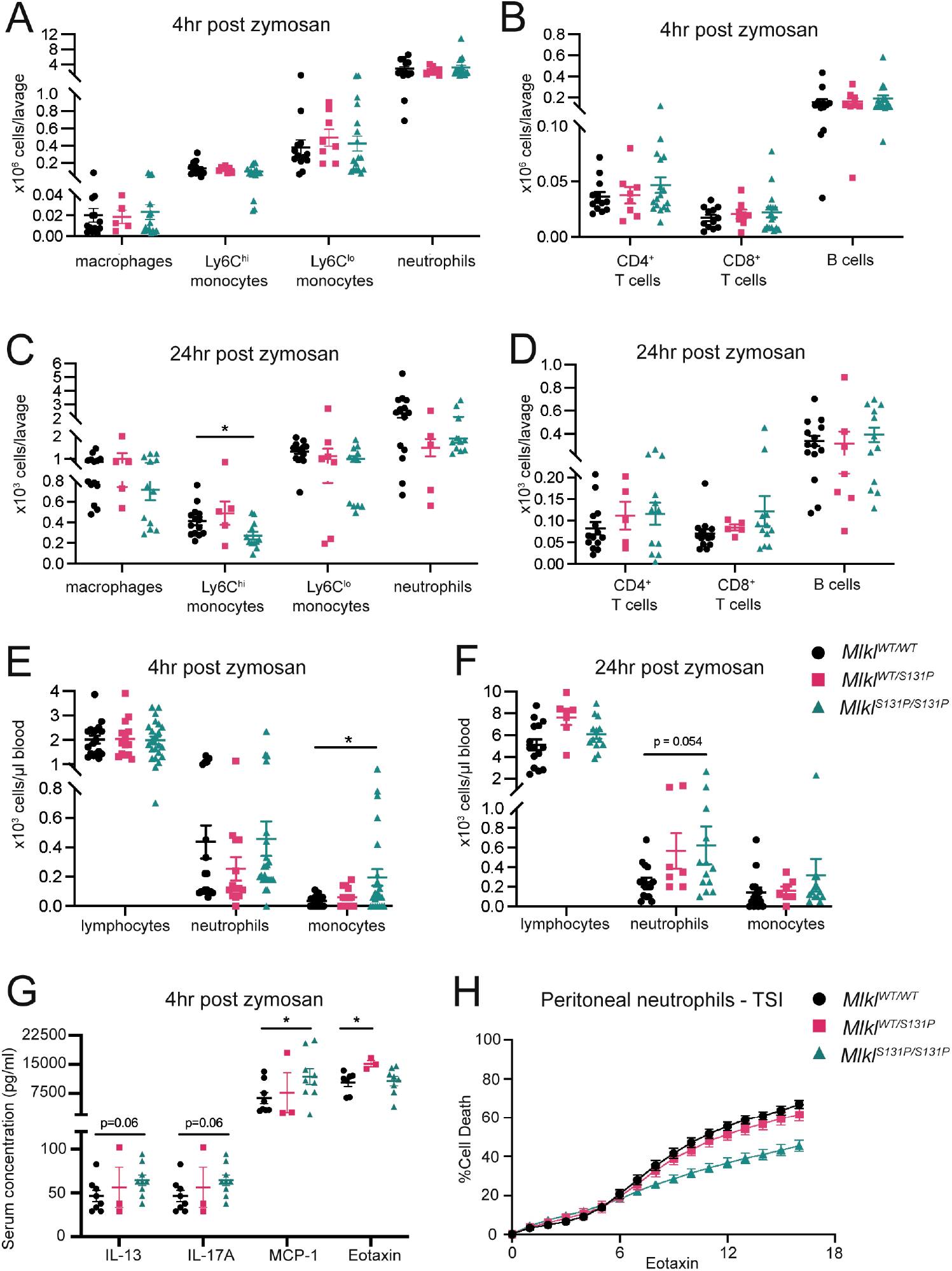
*Mlkl^S131P^* recruited inflammatory neutrophils are less sensitive to TNF-induced necroptosis. **(A-D)** Flow cytometry quantification of peritoneal innate (macrophages, Ly6C^hi^ monocytes, Ly6C^lo^ monocytes and neutrophils) and adaptive (CD4^+^ T cells, CD8^+^ T cells and B cells) immune cells at 4- **(A, B)** or 24- **(C, D)** hours post zymosan injection. ADVIA hematology quantification of circulating immune cells (lymphocytes, neutrophils and monocytes) at 4- **(E)** and 24- **(F)** hours post zymosan injection. **(G)** Multiplex measurement of IL-13, IL-17A, MCP-1 and Eotaxin cytokine levels in peritoneal lavage at 4 hours post-zymosan injection. **(A-G)** Each symbol represents one independent animal, with mice from the 4- or 24-hour timepoint pooled from 3 and 2 independent zymosan experiments respectively. Error bars represent mean ± SEM for *n=3-14* mice as indicated **(A-G)**. Evaluation of induced necroptotic signaling **(H)** in neutrophils recruited and isolated from the peritoneum 4 hours post-zymosan injection. Neutrophils were treated with necroptotic stimulus (TNF, Smac mimetic, IDN-6556; TSI) for 16 hours and cell death was measured every hour by percentage of SYTOX Green positive cells quantified using IncuCyte SX5 live cell imaging. Data were collected from one independent experiment with male and female data pooled, neutrophils isolated from independent mice with mean ± SEM of *n=*6-14 presented. ***p<0.001 calculated using an unpaired, two-tailed Students t-test.

We previously observed that *Mlkl^S131P/S131P^* cells exhibited reduced response to TSI-induced necroptosis. To assess whether this defect in necroptotic function was observed in activated immune cells recruited to sites of inflammation, we isolated neutrophils from the peritoneum at 4 hours post-zymosan injection. Consistent with our previous *in vitro* findings, no differences were observed in the capacity of *Mlkl^S131P/S131P^* inflammatory neutrophils to undergo spontaneous apoptosis **(Supplementary Figure 4Z).** However, the percentage of *Mlkl^S131P/S131P^* inflammatory neutrophils dying following necroptotic stimulation with TSI was decreased in comparison to wild-type neutrophils **(Figure 4H)**.

### *Mlkl^S131P^* homozygotes exhibit reduced capacity to clear *Salmonella* infection

Reductions in the recruitment of Ly6C^hi^ monocytes to sites of inflammation raise important questions as to whether the *Mlkl^S131P^* mutation impacts defense against pathogens. *Salmonella enterica* has long been an important pathogenic selective pressure in humans ^50^. For non-typhoidal *Salmonella* serovars, such as *S. enterica* serovar *Typhimurium*, human infection is typically limited to the gastric mucosa ^51, 52^. In mice and immunocompromised humans, *S. Typhimurium* can cause severe systemic disease after dispersal from the gut by dendritic cells and macrophages to peripheral organs including the spleen and liver ^53^.

*Mlkl^S131P^*homozygote, heterozygote and wild-type mice were infected with a metabolically growth-attenuated *Salmonella typhimurium* strain BRD509 (from here referred to as *Salmonella),* via oral gavage (1 x 10^7^ colony forming units (CFU)). Daily monitoring for the 14 day infection period showed no differences in core body temperature or body mass between genotypes **(Supplementary Figure 5A, B)**. Consistent with normal weights, there were no obvious differences in the integrity of the intestinal epithelial barrier or intestinal monocyte and macrophage populations at infection endpoint **(Figure 5A)**. Despite this, *Mlkl^S131P/S131P^*mice had increased *Salmonella* burden in both the spleen and liver compared to wild-type controls **(Figure 5B, Supplementary Figure 5C, D).** Bacterial colonization in the feces was increased in female *Mlkl^S131P/S131P^* mice relative to wild-type female controls **(Figure 5B, Supplementary Figure 5E)**. In the peripheral blood, *Mlkl^S131P/S131P^*mice had significantly reduced numbers of circulating lymphocytes and monocytes, as well as a trend towards decreased numbers of circulating neutrophils **(Figure 5C, Supplementary Figure 5F-H)**. Infected *Mlkl^S131P/S131P^*mice also exhibited significant decreases in the quantity of splenic Ly6C^hi^ monocytes in comparison to infected wild-type controls **(Figure 5D, E, Supplementary Figure 5I-O)**. At 14 days post-infection, *Mlkl^WT/S131P^* and *Mlkl^S131P/S131P^* mice both had significant increases in plasma concentrations of MCP-1, when compared to wild-type controls **(Figure 5F)**. All other plasma cytokines were equivalent between genotypes at 14 days post-infection **(Figure 5G)**. Splenic (Ly6C^hi^) and circulating monocytes were still significantly decreased in homozygous mice.

**Figure 5.**
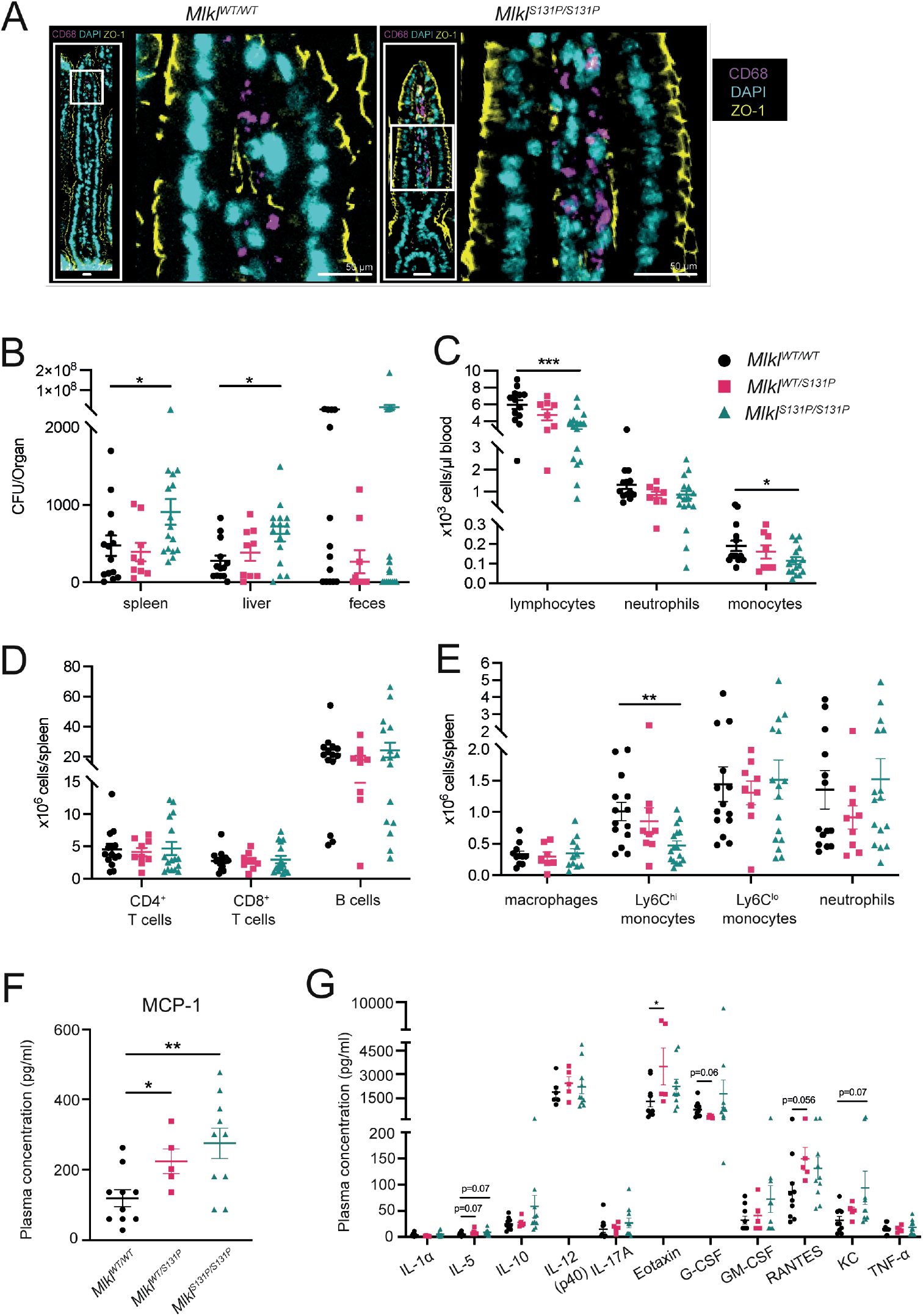
*Mlkl^S131P^* homozygote mice exhibit bacterial clearance defects following oral *Salmonella* infection. **(A)** ZO-1 (yellow) and CD68 (purple) staining in epithelial barrier of intestinal sections taken at 14 days post-*Salmonella* infection. **(B)** Increased bacterial burden observed in spleen and liver, but not feces in *Mlkl^S131P/S131P^* mice at infection endpoint. Quantification of circulating white blood cells (lymphocytes, neutrophils and monocytes) using ADVIA hematology **(C)** and splenic adaptive (CD4^+^ T cells, CD8^+^ T cells and B cells) and innate (macrophages, Ly6C^hi^ monocytes, Ly6C^lo^ monocytes and neutrophils) immune cells using flow cytometry **(D, E)**. **(F, G)** Multiplex measurement of plasma cytokine levels at 14-days post *Salmonella* infection. *Salmonella* infection was performed on 3 independent occasions, with each symbol representing an individual mouse sampled. Error bars represent mean ± SEM for *n=5-16* as indicated. *p<0.05, **p<0.01, ***p<0.001 calculated using a Mann-Whitney test **(B)** or an unpaired, two-tailed Students t-test **(C-G)**.

We did not observe any differences in the kinetics of *Salmonella* induced cell death in *ex vivo* BMDM infection assays, nor inflammasome activation, measured by GSDMD cleavage **(Supplementary Figure 5P, Q)**. Together these data show a hindered pathogen defense in *Mlkl^S131P^* homozygotes accompanied by widespread immunophenotypic deficiencies. Combined, our investigations of *Mlkl^S131P^* homozygotes under challenge highlight a defect in emergency hematopoiesis that manifests in a disruption to integral inflammatory and immune responses. This provides important insights into the potential modulation of immunoinflammatory disorders in *MLKL S132P* carriers.

## DISCUSSION

A high frequency *MLKL S132P* polymorphism present in 2-3% of the global population confers a gain-of-function to MLKL resulting in hematopoietic dysfunction and immune cell defects in a genetically modified mouse model. In human cells, MLKL^S132P^ was resistant to chemical inhibition by necrosulfonamide treatment and endogenous inhibitory phosphorylation at Serine 83. A gain in necroptotic function was also observed when the murine equivalent, *Mlkl^S131P^*, was examined *in situ.* Fibroblasts isolated from *Mlkl^S131P^* homozygotes exhibit RIPK3-independent cell death in the presence of IFN-β, a strong inducer of *Mlkl* gene expression. Under regular culture conditions, MLKL^S131P^ protein levels are reduced relative to MLKL^WT^, manifesting in a reduced capacity for necroptosis in fibroblasts and immune cells stimulated with TSI. While the reduction in MLKL^S132P^ protein levels was also evident in PBMCs isolated from one human individual heterozygous for this polymorphism, our capacity to fully compare between mouse and human systems is significantly limited by the lack of suitable human cell lines that express *MLKL^S132P^* from its endogenous gene locus.

Extensive characterization of the *Mlkl^S131P^* mouse unveiled steady state decreases in the bone marrow pool of inflammatory Ly6C^hi^ monocytes, that were also reflected in reduced numbers at sites of sterile inflammation and bacterial infection. During zymosan-induced peritonitis, increased peripheral monocytes were observed at 4 hours however, Ly6C^hi^ monocytes were significantly reduced in the peritoneum of homozygotes at 24 hours. Under infection challenge, an increased burden of *Salmonella* was present in the spleen and liver of homozygotes, accompanied by significant reductions in the circulating lymphocytes and monocytes. In both cases, monocyte deficits were accompanied by an increase in plasma MCP-1 levels. These deficits are not attributed to any inherent deficits in *Mlkl^S131P^* mouse monocyte/macrophage progenitor cell populations at steady state, nor in their capacity to differentiate and form colonies when measured *ex vivo*. Hematopoietic dysfunction in *Mlkl^S131P^* homozygotes is however evident following radio-ablation, with a severely reduced capacity for stem cells to repopulate *in situ*, or even *Mlkl^WT^* recipients. Hematopoietic stem cells expressing the constitutively active *Mlkl^S131P^* are characterized by increased ROS and Annexin V positivity following irradiation, marking their enhanced propensity for cell death. Taken together, the capacity for monocyte generation, activation and localisation within the *Mlkl^S131P^* mouse under conditions of stress has emerged as the most enticing area for future investigation.

Prior to this work two other single point mutant *Mlkl* mouse models, *Mlkl^D139V^* and *Mlkl^S83G^*, had been reported, both of which exhibited full or partial homozygous postnatal lethality characterized by severe inflammation ^30, 34^. In stark contrast to these models, the *Mlkl^S131P^* homozygotes were born normal and with no signs of inflammatory disease at steady state. However, upon challenge, similarities between *Mlkl^S131P^* and *Mlkl^D139V^* mice were revealed. Hematopoietic dysfunction is observed following myelosuppressive irradiation in homozygous or heterozygous mice harboring the *Mlkl^S131P^* or *Mlkl^D139V^* mice, respectively ^34^.

Despite the ostensible basal state phenotypes of *Mlkl^D139V^* and recently described *Mlkl^S83G^* homozygotes differing from the *Mlkl^S131P^* homozygotes^30, 34^, there were similarities observed at the molecular level. As for MLKL^S131P^, RIPK3-independent gain-of-function was reported for MLKL^D139V^, but not MLKL^S83G 30, 34^. This major similarity in molecular function is not unexpected considering the proximity of the D139 and S131 residues within MLKL. Constitutive activity of the D139V brace helix mutants is restrained by cellular mechanisms that limited protein level, with decreases in endogenous MLKL observed in cells generated from *Mlkl^D139V^* homozygotes. Interestingly, ubiquitin-mediated targeting of MLKL to the lysosome was identified to be mechanistically involved in the restraint of necroptosis in cells expressing MLKL^D139V^, and as a means of clearing endosomal *Listeria monocytogenes* and *Yersinia enterocolita* ^34, 54, 55^. Exploring if the reduction in cellular MLKL levels we observed in the PBMCs of a human carrier of the *MLKL^S132P^*polymorphism, or cells isolated from the *MLKL^S131P^* mouse, is likewise mediated by ubiquitination and lysosomal targeting will be an important next step, particularly as it relates to the clearance of these intracellular bacteria from human cells. Ablation of the inhibitory phosphorylation site in *Mlkl^S83G^* mice similarly subjects MLKL to downregulation in mouse cells ^30^. In sum, our analysis of MLKL^S131P^ further highlights the importance of cellular mechanisms that clear activated MLKL from cells below a threshold to reduce aberrant necroptotic cell death.

The finding that MLKL^S132P^ retains necroptotic killing activity despite simulated inhibitory phosphorylation at Ser83 in human cells is highly notable. The inflammatory phenotypes that result from homozygosity of the phosphoablating(S83G) mutation provides insight into the potential disease development that may occur in carriers of the *MLKL S132P* polymorphism under environmental and cellular scenarios where Ser83 inhibitory phosphorylation is deployed. Identifying the kinase(s) responsible for phosphorylation at MLKL Ser83 and understanding which cell types are primed to deploy this kinase(s) is an important next step in determining if this polymorphism promotes clinically relevant changes to homeostasis or disease outcomes in humans.

Evidence for positive selection has been found for over 300 immune-related gene loci and many of these have been found to be associated with the incidence of autoimmune and autoinflammatory disease in modern humans ^56, 57^. Many of these variants have also been mechanistically linked to pathogen defense (Karlsson et al., 2014, Ramos et al., 2015), with pathogenic microbes a major driver of genetic selection over human history. While a diminished capacity for mice to clear disseminated *Salmonella* would argue against the hypothesis that the *MLKL S132P* polymorphism has been positively selected for in human populations, it is important to note that only 1.4 % of human carriers are homozygotes ^40^. *MLKL S132P* heterozygosity is by far the most prevalent, and evolutionarily relevant, human scenario. *Mlkl^S131P^* heterozygote mice displayed gene dosage phenotypes consistent with homozygotes, with one exception. Following sublethal myelosuppressive irradiation, a gain in bone marrow hematopoietic stem cell numbers in heterozygotes was observed at day 21 whilst a catastrophic drop occurred in homozygotes. This increased stem cell capacity does not persist long-term. In competitive bone marrow transplants, *Mlkl^WT/S131P^* stem cells compete similarly to *Mlkl^WT/WT^* stem cells at 6 weeks post-transplantation. However, the increased fitness conferred to heterozygous mice at this early timepoint provides intriguing insights into other selective pressures that may have promoted accumulation of this polymorphism in humans ^58^. Sepsis, where emergency hematopoiesis is an essential determinant in survival, is an interesting and highly evolutionarily important avenue of exploration for the study of *MLKL S132P* polymorphism frequency ^59, 60^. Since nutrition is another important driver of genetic adaptation in humans, exploring the role of *MLKL S132P* in metabolic disease is also of interest. An important precedent for this is a recent report of an association between human RIPK1 promoter polymorphisms and diet-induced obesity ^61^. The potential for negative selection of this polymorphism over time is also an important avenue of exploration in light of the recent discovery of a common *TYK2* variant and its role in enhancing susceptibility to severe infection by historically important human pathogen mycobacterium tuberculosis ^62, 63^.

By many, MLKL is viewed as a potential therapeutic target for drug discovery due to its established involvement in multiple human diseases, especially inflammatory pathologies ^34, 64, 65, 66^. To date no human clinical trials have been conducted on MLKL-targeted small molecules, although several inhibitors have been reported in the literature, including human MLKL inhibitor necrosulfonamide (NSA), which all function by targeting Cys86 ^16, 67^. Intriguingly, we show that in the presence of NSA human MLKL^S132P^ protein displays a gain-of-function that results in non-inhibitable cell death. While NSA has been a fundamental tool for *in vitro* studies of necroptosis, our findings raise interesting questions about the suitability of Cys86-targetted MLKL inhibitors for the 2-3% of the population that carry the *MLKL S132P* polymorphism.

## METHODS

### Patient Recruitment, Ethics and Informed consent

Patients and their relatives were recruited from the Department of Clinical Immunology and Allergy, Royal Melbourne Hospital, Victoria, Australia and the Centre for Personalized Immunology, Australian National University, Canberra, Australia. Unrelated, age and sex matched ‘healthy’ controls that did not carry the *MLKL p.Ser132Pro* polymorphism were recruited via the Volunteer Blood Donor Registry, Parkville. Informed consent was obtained from all participants for genomic analysis and immunological studies. All procedures were performed and are reported here with the approval of human ethics review boards of all Institutes that participated in human genetics studies; Australian National University, The Walter and Eliza Hall Institute of Medical Research (approved projects 2009.162, 10/02) and with the 1964 Helsinki declaration and its later amendments or comparable ethical standards.

### Genomic analysis

Whole exome sequencing was performed by the Canberra Clinical Genomics service. Libraries were prepared and enriched using the SureSelect Clinical Research Exome v2 kit (Agilent Technologies), and targeted regions were sequenced using an Illumina sequencing system with 100bp paired-end reads, with an average depth of >35. Reads were aligned to the human genome reference sequence (GRCh38) using the Burrows-Wheeler Aligner (BWA-MEM), and variant calls made using the Genomic Analysis Tool Kit (GATK).

### Animal ethics

All mice were housed at the Walter and Eliza Hall Institute of Medical Research (WEHI), Australia. This facility is a temperature and humidity controlled specific pathogen free facility with a 12h:12h day night cycle. All experiments were approved by the WEHI Animal Ethics Committee in accordance with the Prevention of Cruelty to Animals Act (1986) and the Australian National Health and Medical Research Council Code of Practice for the Care and Use of Animals for Scientific Purposes (1997).

### Mice

Mice were generated on a C57BL/6J background. The *p.Ser131Pro* mutation in the *Mlkl* gene (*Mlkl^S131P^*) was generated using CRISPR/Cas9 by the Melbourne Advanced Genome Editing Centre (MAGEC) laboratory at WEHI, following the same methodology as previously described^34, 68^. To introduce a proline-coding mutation in place of Ser131 within the *Mlkl* gene, an sgRNA of the sequence CTGTCGATCTTCCTGCTGCC was used to create double stranded breaks within the *Mlkl* locus and initiate homologous recombination.

An oligo donor (TGTTGCTGCTGCTTCAGGTTTATCATTGGAATACCGTTTCAGATGTCAGCCAGCCAGCACCATGGCAGCAGGAAGATCGACAGGATGCAGAGGAAGACGGgtgagtctcccaaagactgg ga) was subsequently used to introduce the S131P mutation. Genotyping of mice was completed by Transnetyx using custom probe Mlkl-8 MUT (Forward Primer: CTGCTTCAGGTTTATCATTGGAATACC, Reverse Primer: TCTGCATCCTGTCGATCTTCCT).

### Reagents

Antibodies; Rat anti-mMLKL 8F6^34^(1:2000), rat anti-mMLKL 5A6^69^ (1:1000; available from Merck-Millipore as MABC1634), rat anti-hMLKL 10C2^22^ (1:1000; available from Merck-Millipore as MABC1635), rat anti-hMLKL 7G2^22^ (1:1000; available from Merck-Millipore as MABC1636), rat anti-hRIPK3 1H2^4^ (1:1000; available from Merck-Millipore as MABC1640) were produced in-house. Mouse anti-actin (A-1978; 1:5000) was purchased from Sigma-Aldrich, rabbit anti-GAPDH (#2118; 1:2000-5000) was purchased from Cell Signalling Technology, rabbit anti-VDAC (AB10527; 1:10000) was purchased from Millipore, rabbit anti-human pMLKL (EPR9514; 1:1000-3000) was purchased from Abcam, and rabbit anti-mouse pMLKL (D6E3G; 1:1000) was purchased from CST. Cell treatments were completed with agonists/antagonists at the following concentrations: 100 ng/ml recombinant hTNF-Fc (produced in house as in ^70^), 500 nM Smac mimetic Compound A (provided by Tetralogic Pharmaceuticals; as in ^71^, 5 µM Pan-caspase inhibitor IDN-6556 (provided by Tetralogic Pharmaceuticals), 1 µM necrosulfonamide (NSA; Merck #480073), 5 µM necrostatin (Nec-1s; Merk #504297), 1 µM GSK′872 (SynKinase #SYN-5481), 10-20 ng/ml lipopolysaccharide (LPS; Sigma #L2630), 25 µg/ml polyinosinic:polycytidylic (Poly I:C; Novus), 200 nM MG132 (Merck #474790), 2 nM PS341 (Sigma #504314), and 30 ng/ml mouse IFNβ (R&D Systems #8234-MB-010)

### Generation of cell lines

Mutations were introduced into a human MLKL DNA template (from DNA2.0, CA) using oligonucleotide-directed PCR and sub-cloned into the pF TRE3G PGK puro vector^18^. Vector DNA was co-transfected into HEK293T cells with pVSVg and pCMV δR8.2 helper plasmids to generate lentiviral particles. U937 (WT and *MLKL^-/-^*) and HT29 (WT and *MLKL^-/-^*) were then stably transduced with exogenous human MLKL ligated into pFTRE3G. Successfully transduced cells were selected using puromycin (2.5µg/mL; StemCell Technologies) using established procedures^18, 27, 46^. The following oligonucleotides were used for the assembly of constructs:

hMLKL^S132P^ fwd; 5′ GCCAAGGAGCGCCCTGGGCACAG3′

hMLKL^S132P^ rev; 5′ CTGTGCCCAGGGCGCTCCTTGGC 3′

hMLKL^TSEE^ fwd; 5′ GAGGAAAACACAGGAGGAAATGAGTTTGGGAAC 3′

hMLKL^TSEE^ rev; 5′ GTTCCCAAACTCATTTCCTCCTGTGTTTCCTC 3′

hMLKL^S83A^ fwd; 5′ GTTCAGCAATAGAGCCAATATCTGCAG 3′

hMLKL^S83A^ rev; 5′ CCTGCAGATATTGGCTCTATTGCTGAAC 3′

hMLKL^S83D^ fwd; 5′ GAAAAGTTCAGCAATAGAGACAATATCTGCAGGTTTC 3′

hMLKL^S83D^ rev; 5′ GAAACCTGCAGATATTGTCTCTATTGCTGAACTTTTC 3′

hMLKL^S83C^ fwd; 5’ GTTCAGCAATAGATgCAATATCTGCAGG 3’

hMLKL^S83C^ rev; 5’ CCTGCAGATATTGcATCTATTGCTGAAC 3’

hMLKL^R82S^ fwd; 5’ GAAAAGTTCAGCAATAGCTCCAATATCTGCAG 3’

hMLKL^R82S^ rev; 5’ CTGCAGATATTGGAGCTATTGCTGAACTTTTC 3’

Primary mouse dermal fibroblasts (MDFs) were prepared from skin taken from the head and body of 1 day old mice. These MDFs were immortalised by stable lentiviral transduction with SV40 large T antigen. Bone marrow derived macrophages were generated from the long bones of adult mice and grown for 7 days in DMEM supplemented with 15% L929 cell supernatant.

Human blood (patient and healthy donor) was collected by collaborators via venipuncture at the Royal Melbourne Hospital. Collected blood was diluted with PBS and layered on an equal volume of Histopaque (density 1.077 g/ml) and centrifuged for 30 minutes, 700 x g at 20 °C. The layer containing peripheral blood mononuclear cells (PBMCs) was harvested and washed with PBS, then frozen in FCS + 10 % DMSO and stored in liquid nitrogen.

### Culture of cell lines

Primary MDFs, immortalised MDFs and HT29s were cultured in DMEM + 8% FCS. BMDMs were cultured in DMEM + 15% FCS + 20% L929. U937 and PBMCs were cultured in RPMI + 8% FCS. All cell lines were grown at 37 °C and 10% (v/v) CO_2_.

### Western blot

U937 cells were seeded into 48-well plates at 60,000 cells/well and induced for 3 hours with doxycycline (20 ng/mL, 100 ng/mL or 500 ng/mL) to stimulate MLKL expression. HT29 cells were seeded into 48-well plates at 45,000 cells/well and following 12-14 hours of cell adherence, cells were induced overnight with doxycycline (20 ng/mL, 100 ng/mL or 500 ng/mL) for stimulation of MLKL expression. BMDMs were plated at 400,000 cells/well in a 24-well plate and MDFs (primary and immortalised) were plated at 30,000 cells/well in a 48-well plate. Cells were stimulated as indicated at described for 6 hours, except for BMDMs stimulated with LPS for 2 hours before addition of Smac mimetic Compound A for a further 4 hours. Human primary PBMCs were plated at 45,000 cells/well and stimulated for 4 hours. All cells were harvested in 2 x SDS Laemmli’s lysis buffer, boiled at 100 °C for 10-15 min, and then resolved by 4-15% Tris-Glycine SDS-PAGE (Bio-Rad). Proteins were transferred to nitrocellulose or PVDF membrane and probed with antibodies as indicated.

### IncuCyte analysis

Primary and immortalised MDFs were plated at 8,000 cells per well in a 96-well plate. BMDMs were plated at 150,000 cells per well on day 6 of culture in a 48-well plate. MDFs and BMDMs were left to settle overnight before stimulation. The next day MDFs and BMDMs were stimulated in culture media supplemented with propidium iodide. HT29 cells were plated at 45,000 cells per well in a 48-well plate and left to settle for 6 hours before overnight doxycycline pre-stimulation (20ng/mL, 100ng/mL or 500ng/mL). HT29 cells were stimulated in Phenol Red-free media supplemented with 2% FCS, 1mM Na pyruvate, 1mM L-GlutaMAX, SYTOX Green (Invitrogen, S7020) and either DRAQ5 (Thermofisher, #62251) (MLKL KO HT29) or SPY_620 (Spirochrome, SC401) (WT HT29).

U937 cells were plated at 60,000 cells per well in a 48-well plate and were induced with doxycycline (20ng/mL, 100ng/mL or 500ng/mL) for 3 hours. Cells were then stimulated in Phenol Red-free media supplemented with 2% FCS, 1mM Na pyruvate, 1mM L-GlutaMAX, SYTOX Green and either DRAQ5 (MLKL KO HT29) or SPY_620 (WT HT29).

Neutrophils isolated from the peritoneum at 4 hours post-zymosan injection were counted and plated at 60,000 cells/well in a 48-well plate. Plating media (RPMI + 8 % FCS) was supplemented with SYTOX Green and DRAQ5 dyes.

Images were taken every hour using IncuCyte SX5 or S3 imaging and cell death was quantified by number of dead cells (SYTOX Green or propidium iodide positive). Percentage values were quantified by number of dead cells out of total live cell number (DRAQ5 or SPY620 positive).

### TNF ELISA

100,000 cells were stimulated with LPS (10ng/mL) or Poly I:C (2.5µg/mL) for 3 hours. PBMC supernatant cytokine content was measured by ELISA (R&D: STA00C) according to the manufacturer’s instructions. The measurements were performed in technical triplicates.

### Mouse histopathology

7–9-month-old mice were euthanized by CO_2_ and fixed in 10 % buffered formalin. For the full body, 5-µm sagittal sections were taken at 300-µm intervals of all organs. Histopathologists Aira Nuguid and Tina Cardamome at the Australian Phenomics Network, Melbourne completed thorough examination of these sections.

### Haematological analysis

Cardiac, submandibular or retro-orbital blood collected from mice at steady state (8-52 weeks old) or following challenge was placed into EDTA coated tubes. Blood cells were left undiluted or diluted 2- to 11-fold in DPBS for automated blood cell quantification using an ADVIA 2120i haematological analyser on the same day as harvest.

### Cytokine quantification

All plasma was collected by centrifugation (10,000 g, 5 min) and stored at -80 °C. Lavage fluid and plasma cytokine quantities were measured by Bioplex Pro mouse cytokine 23-plex assay (Bio-Rad #M60009RDPD) according to manufacturer’s instructions. When samples were denoted as ‘<OOR’, below reference range, for a particular cytokine they were assigned the lowest recorded value for that cytokine across all samples.

### Colony forming assays

Single-cell suspensions from adult bone marrow were prepared in balanced salt soluiton (0.15 M NaCl, 4 mM KCl, 2mM CaCl_2_, 1mM MgSO_4_, 1mM KH_2_PO_4_, 0.8 mM K_2_HPO_4_, and 15 mM N-2-hydroxyethylpiperazine-N’2-ethanesulfonic acid supplemented with 2% [v/v] bovine calf plasma). Clonal analysis of bone marrow cells (2.5 x 10^4^) was performed in 1 mL semisolid agar cultures of 0.3% agar in DMEM containing 20% newborn calf plasma, stem cell factor (SCF; 100 ng/mL; in-house), erythropoietin (EPO; 2 U/mL; Janssen), interleukin-3 (IL-3; 10 ng/mL; in-house), G-CSF (10^3^ U/mL; PeproTech), granulocyte-macrophage colony stimulating factor (M-CSF; 10^3^ U/mL; in-house). Cultures were incubated at 37 °C for 7 days in a fully humidified atmosphere of 10% CO2 in air, then fixed, dried onto glass slides, and stained for acetylcholinesterase, luxol fast blue, haematoxylin, and the number and type of colonies were determined, blinded.

### Mouse model of *Salmonella* infection

Mice used in this experiment were a mix of littermates and non-littermates aged 6-12 weeks, and wild-type mice that were littermates behaved equivalently to non-littermates. Mice were infected with *Salmonella enterica* serovar Typhimurium strain BRD509 ^72^ at 10^7^ colony forming units (CFU) by oral gavage. Mice were harvested 14 days post-infection. Cardiac bleeds were taken, and blood populations analysed using an ADVIA hematology analyser. Liver, spleen, and faeces were harvested for enumeration of viable bacteria on nutrient agar. Organs from infected mice were weighed and homogenised in 2 mL (spleens), 5 mL (livers) or 1 mL (faeces) of PBS. Homogenates were serially diluted (in duplicate) in PBS and 10 µl drops plated out in duplicate onto LB agar (+ streptomycin) and incubated overnight at 37 °C. CFU/mL was calculated per organ for each mouse and then standardised to CFU/organ based upon organ weight. A small portion (1/3) of the spleen was processed for flow cytometry analysis.

### *In vitro Salmonella* infection

*In vitro* infection of BMDMs with *Salmonella enterica* serovar Typhimurium strain SL1344 were performed as previously reported (Doerflinger et al., 2020). BMDMs on day 6 of differentiation were plated at 4 x 10^5^ cells/well in a 6-well plate for western blot analyses or 5 x 10^4^ cells/well in a 96-well plate for LDH assays. Cells were infected at MOI:25 for western blot or MOI:10 or 50 for LDH assays in antibiotic-free DMEM for denoted incubation times. For all experimental analyses, following 30 minute incubation, cells were washed and replaced in DMEM media supplemented with 50 µg/ml gentamycin to ensure growth inhibition of extracellular bacteria. BMDM cell death levels were measured as a percentage of LDH release using the Promega CytoTox 96 Non-Radioactive Cytotoxicity Assay (G1780), according to manufacturer’s instructions.

### Sublethal irradiation

Mice used in this experiment were a mix of littermates and non-littermates aged 7-17 weeks, and wild-type mice that were littermates behaved equivalently to non-littermates. Mice were irradiated with a 5.5 Gy sub-lethal dose of γ-irradiation and received neomycin (2 mg/mL) in the drinking water for 3 weeks. At 4 days post irradiation, a retro-orbital bleed was analysed via ADVIA hematology to confirm successful irradiation. Mice had submandibular bleeds analysed by ADVIA hematology and had long bones harvested for flow cytometry analysis at either 7, 14 or 21 days post-irradiation to assess stem cell capacity.

### Hematopoietic stem cell transplants

Donor bone marrow were injected intravenously into recipient *C57BL/6-CD45^ly5.1^* mice following 11 Gy of γ-irradiation split over two equal doses. Recipient mice received neomycin (2 mg/mL) in the drinking water for 3 weeks. Long term capacity of stem cells was assessed by flow cytometric analysis of donor contribution to recipient mouse peripheral blood at 6 weeks.

### Zymosan-induced peritonitis

Mice used in this experiment were a mix of littermates and non-littermates aged 7-17 weeks, and wild-type mice that were littermates behaved equivalently to non-littermates. 1 mg of zymosan A from *Saccharomyces cerevisiae* (Sigma-Aldrich) was intra-peritoneally injected into mice to induce sterile peritonitis. After 4 or 24 hours, mice were euthanized, cardiac bled and bone marrow collected. The peritoneal cavity was washed with 1 ml of cold PBS and cells within the lavage fluid were collected by centrifugation (300 x g, 5 minutes). Bone marrow and peritoneum immune cells were quantified by flow cytometry.

### Flow cytometry

To analyse the innate and adaptive immune cells in peripheral blood, inguinal lymph nodes, spleen and bone marrow, isolated single cell suspensions were incubated with a combination of the following antibodies: CD4-BV421, CD8-PECy7, CD19-PerCPCy5.5, CD11b-BV510 or BV786, GR1-PE, CD45-Alexa700, Ly6G-V450 and Ly6C-APCCy7.

Splenic immune cells were analysed at 14-day post *Salmonella* infection and incubated with a combination of the following antibodies: CD4-BV421, CD8-PeCy7, CD19-PerCPCy5.5, CD11b-BV510 or BV786, CD64-PE, CD45-Alexa700, Ly6G-V450 and Ly6C-APCCy7.

To analyse the contribution of donor and competitor cells in transplanted recipients, blood cells were incubated with either CD41-APC (ThermoFisher) and Ter119-PE or Ly5.1-A700, Ly5.2-PE-Cy7, CD4-PE, CD8-PE, B220-BV650, Ly6C-APCCy7, Mac1-PerCPCy5.5. To analyse stem- and progenitor-cell compartment following sub-lethal irradiation, bone marrow cells were incubated with cKit-PerCPe710/PerCPCy5.5 (ThermoFisher), CD48-PECy7, CD150-A647, Sca1-APCCy7, B220-PE, CD19-PE, CD4-PE, CD8-PE, Gr1-PE. To analyse the relative percentages of stem and progenitor cells at steady state, bone marrow cells were incubated with cKit-PerCPCy5.5, Sca1-A594, CD150-BV421, CD105-PE, FcγRII-PECy7, and lineage markers (B220, CD19, CD4, CD8, GR1, Ly6G, Ter119)-A700. Finally, fluorogold (AAT Bioquest Cat#17514) was added for dead cell detection where appropriate. For detection of Annexin V at 21 days post-irradiation, bone marrow was incubated with Annexin V for 30 minutes.

Peritoneal and bone marrow immune cells at 4- or 24-hours post-zymosan injection were incubated with CD45-Alexa700, CD64-BV650, Ly6G-PE, Ly6C-APCCy7, F4/80-PerCPCy5.5, CD11b-BV510, CD8-PECy7, CD4-FITC, B220-APC and FC-blocker. Finally, propidium iodide (2 µg/mL, Sigma-Aldrich) was added for dead cell detection.

All cells were analysed on the Aurora Cytex flow cytometer. With exception of zymosan induced peritonitis experiments that used the Aurora Cytex automated volume calculator, cells were mixed with counting beads to quantify absolute cell numbers. Except where denoted, all flow cytometry antibodies were obtained from BD Biosciences.

### Reactive oxygen species (ROS) detection

ROS was detected by mixing Chloromethyl-H_2_DCFDA dye (1µM; Invitrogen, #C6827) with bone marrow harvested from mice 21 days post-irradiation. Following a 30 minute incubation at 37 °C, loading buffer was removed and cells were placed into StemPro-34 plasma free medium (Thermofisher, #10639011) for 15 minute chase period. Cells were analysed using Aurora Cytex flow cytometer.

### Liposome dye release assays

Recombinant full-length human MLKL (residue 2-471) proteins were expressed in *Sf21* insect cells using bacmids prepared in DH10MultiBac *E.coli* (ATG Biosynthetics) from pFastBac Htb vectors using established procedures^73^. Briefly, full-length GST-tagged human MLKL proteins were purified using glutathione agarose resin (UBPBio) ^46^ followed by size-exclusion chromatography using HiLoad 16/600 Superdex 200 pg column (Cytivia). Fractions corresponding to full-length human MLKL tetramers (elution volume 55-63 ml) were pooled for liposome assays. Liposomes (100 nm diameter) with a plasma membrane-like lipid mix (20% POPE, 40% POPC, 10% PI/PI(4,5)P_2_, 20% POPS, 10% POPG) filled with 50 mM 5(6)-Carboxyfluorescein dye (Sigma) were prepared as previously described^29^. Recombinant human MLKL protein was diluted to 1 µM (2 x desired final concentration) in LUV buffer (10 mM HEPES pH 7.5, 135 mM KCl) and aliquoted into a 96 well flat-bottom plate (ThermoFisher Scientific). Liposomes were purified from excess dye using a PD-10 desalting column (Cytiva) and diluted to 20 µM in LUV buffer. Immediately following addition of the liposomes to the plate (1:1 ratio liposomes:protein) fluorescence (485nm excitation, 535nm emission) was measured every 2 minutes for 60 minutes total on the CLARIOstar plate reader (BMG Labtech). Baseline measurements were determined by incubation of liposomes with LUV buffer alone. All assays were performed in triplicate. Data plotted as mean ± SD of one independent repeat that is representative of three independent assays.

### Statistical analyses

All data points signify independent experimental repeats or biologically independent data points. All *p* values were calculated in Prism using the statistical test identified in figure legends. Asterisks signify that *p* ≤ 0.05 (*), *p* ≤ 0.01(**) or *p* ≤ 0.001 (***).

## ACKNOWLEDGEMENTS

We thank all the following people for their technical assistance; Aira Nuguid and Tina Cardamone (Phenomics Australia Histopathology and Slide Scanning Service-The University of Melbourne). WEHI Cytometry Facility, WEHI Antibody Facility, WEHI Centre for Dynamic Imaging, WEHI Bioservices, Cheree Fitzgibbon (WEHI), Jacinta Hansen (WEHI) and Matthew Cook (ANU). The generation of *Mlkl^S131P^* mice by CRISPR/Cas9 gene editing was performed by Andrew Kueh and Marco Herold (WEHI MAGEC laboratory) supported by the Australian Phenomics Network (APN) and the Australian Government through the National Collaborative Research Infrastructure Strategy (NCRIS) program. We thank Warren Alexander and Melanie Bahlo for the provision of important resources and expertise. We thank Michael Hildebrand and Tom Witkowski from Epilepsy Research Centre, Department of Medicine, Austin Health for assistance with Sanger sequencing. We are grateful to the National Health and Medical Research Council for fellowship (J.M.H., 1142669; A.L.S., 2002965; J.M.M., 1172929; J.S., 1107149), grant (J.M.M., 1105023; K.R.M., 1092602; J.S., 1105023; J.M.H., 2011584) and infrastructure (IRIISS 9000719); Arthritis Australia support to K.R.M; K.E.L funding by Future Fellowships from the ARC (FT19010266). We acknowledge scholarship support for S.E.G, Y.M, D.F and A.V.J (Australian Government Research Training Program Stipend Scholarships), S.E.G (Wendy Dowsett Scholarship), S.C (Walter and Eliza Hall Handman PhD Scholarship). Victorian State Government Operational Infrastructure Support Scheme.

## AUTHOR INFORMATION

**The Walter and Eliza Hall Institute of Medical Research, Parkville, VIC, 3052, Australia**

Sarah E. Garnish, Katherine R. Martin, Maria Kauppi, Victoria Jackson, Shene Chiou,Yanxiang Meng, Daniel Frank, Emma C. Tovey Crutchfield, Komal M. Patel, Annette V.Jacobsen, Georgia K. Atkin-Smith, Ladina Di Rago, Marcel Doerflinger, Christopher R. Horne, Cathrine Hall, Samuel N. Young, Ian P. Wicks, Ashley P. Ng, Charlotte Slade, Andre L. Samson, John Silke, James M. Murphy and Joanne M. Hildebrand

**Department of Medical Biology, University of Melbourne, Parkville, VIC, 3052, Australia**

Sarah E. Garnish, Katherine R. Martin, Maria Kauppi, Shene Chiou,Yanxiang Meng, Komal M. Patel, Annette V.Jacobsen, Georgia K. Atkin-Smith, Ladina Di Rago, Marcel Doerflinger, Christopher R. Horne, Ian P. Wicks, Charlotte Slade, Andre L. Samson, John Silke, James M. Murphy and Joanne M. Hildebrand

**Centre for Innate Immunity and Infectious Diseases, Hudson Institute of Medical Research, Clayton, VIC, 3168, Australia**

Rebecca Ambrose, Vik Ven Eng and Jaclyn S. Pearson

**Department of Molecular and Translational Science, Monash University, Clayton, VIC, 3168, Australia**

Rebecca Ambrose, Vik Ven Eng, Kate E. Lawlor and Jaclyn S. Pearson

**Department of Microbiology, Monash University, Clayton, VIC, 3168, Australia**

Jaclyn S. Pearson

**University of Melbourne, Faculty of Medicine, Dentistry and Health Sciences, Parkville, 3052, Australia**

Emma C. Tovey-Crutchfield

**Department of Immunology and Infection, John Curtin School of Medical Research, Australian National University, ACT, Australia**

Vicki Athanasopoulos and Carola Vinuesa

**Institute of Virology, Technical University of Munich/Helmholtz Munich, Munich, Germany**

Gregor Ebert

**Clinical Haematology Department, The Royal Melbourne Hospital and Peter MacCallum Cancer Centre, Parkville, 3052, Australia**

Ashley P. Ng

**Department of Clinical Immunology and Allergy, Royal Melbourne Hospital, Parkville, VIC, 3052, Australia**

Charlotte A. Slade

### Contributions

Conceptualization: S.E.G., J.S., J.M.M., J.M.H. Methodology: S.E.G., K.R.M., M.K., V.J., R.A., V.E., S.C., Y.M., D.F., E.C.T., K.M.P., A.V.J., G.K.A., L.D., M.D., C.R.H., C.H., S.N.Y., G.E., A.P.N., J.S.P., A.L.S., K.E.L., J.M.H. Resources: K.R.M., M.K., V.A., C.V., C.S., J.S.P., J.S., J.M.M., J.M.H. Supervision: K.M., M.K., J.S., J.M.M., J.M.H. Funding acquisition: J.S., J.M.M., J.M.H. S.E.G and J.M.H co-wrote the paper with input from authors.

### Data Availability

The biological tools generated for MLKL during this study are available from the corresponding authors on reasonable request.

## ETHICS DECLARATIONS

Competing interests: S.E.G, K.M.P, A.L.S, C.R.H, S.N.Y, J.S, J.M.M and J.M.H contribute, or have contributed, to a project developing necroptosis inhibitors in collaboration with Anaxis Pty Ltd. K.R.M received funding from CSL Pty Ltd. The remaining authors declare no competing interests.

## SUPPLEMENTARY FIGURES

**Supplementary Figure 1.**
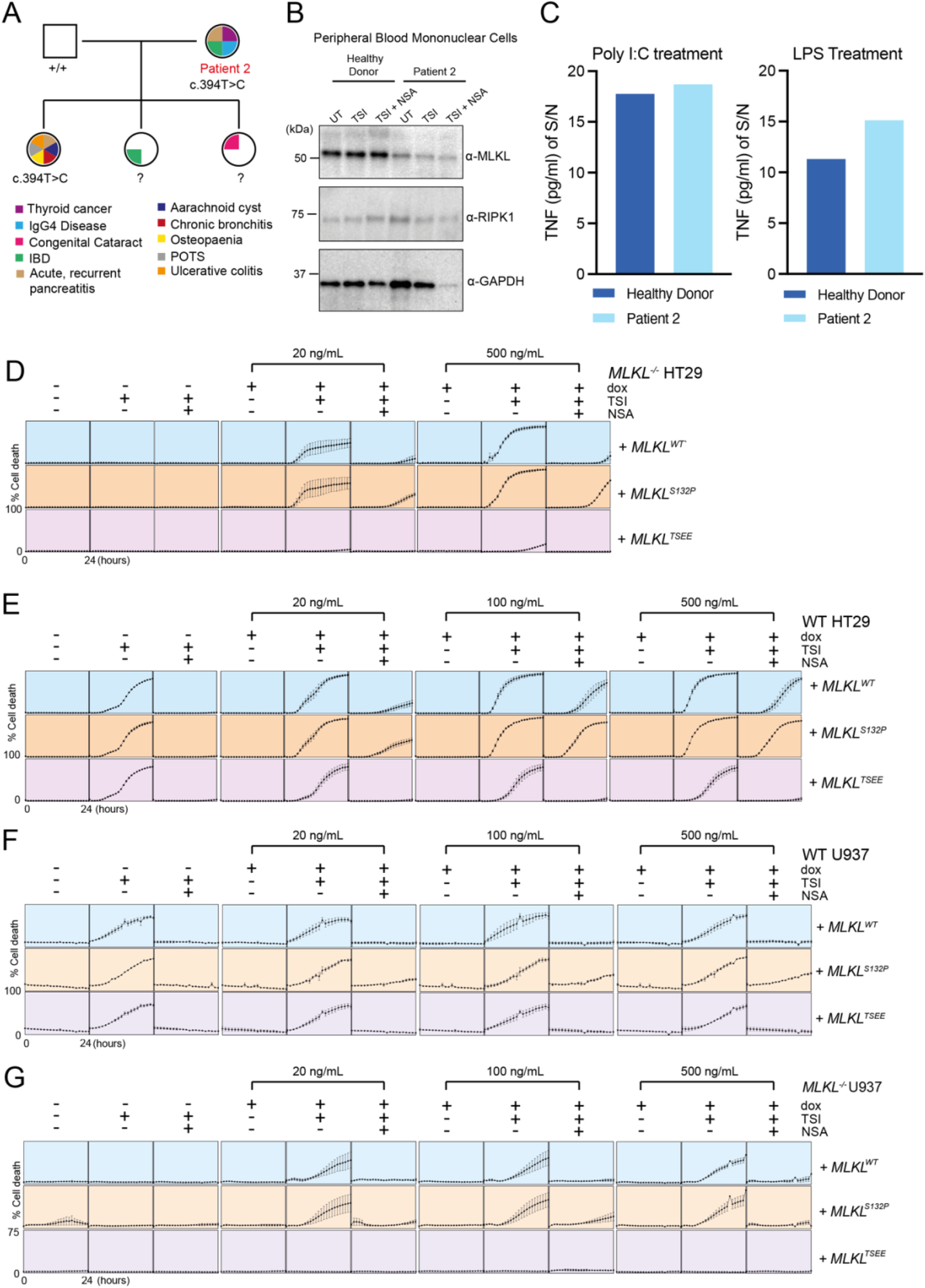
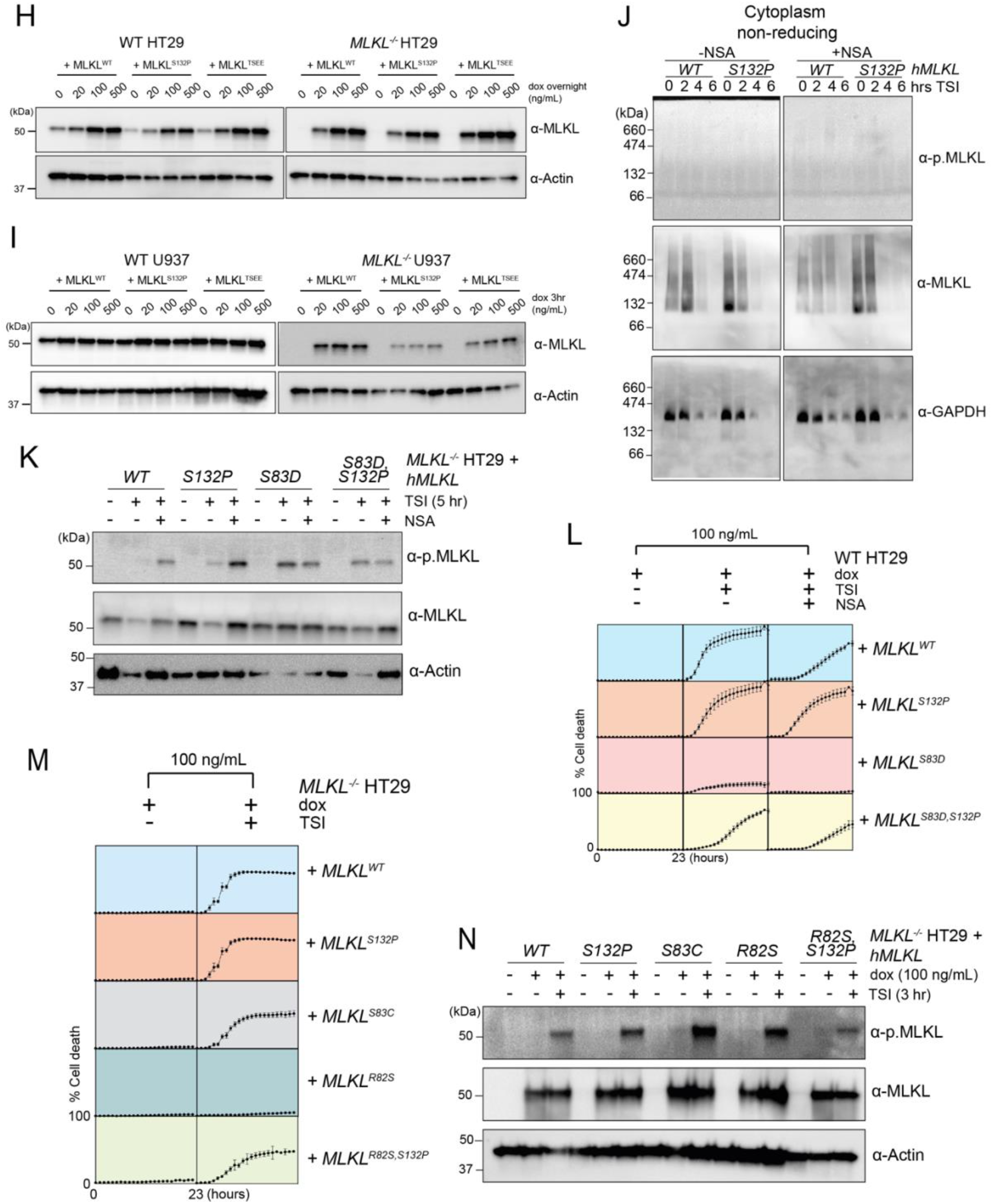
MLKL^S132P^ is less sensitive to inhibition by necrosulfonamide. (**A)** Family pedigree of patient 2, identified from the Australian registry of patients suffering from immune related disease. Known diagnoses for family members are indicated. (**B)** Peripheral blood mononuclear cells (PBMCs) isolated from patient 2 and an aged matched healthy donor control were stimulated as indicated for 4 hours for western blot analysis. (**C)** ELISA measurement of supernatant TNF in PBMCs stimulated with LPS or Poly I:C for 5 hours. Mean of technical triplicates presented. **(D-G)** Evaluation of necroptotic signaling by MLKL^WT^, MLKL^S132P^ and MLKL^TSEE^ in *MLKL^-/-^* **(D)** or WT **(E)** HT29 cells and WT **(F)** or *MLKL^-/-^* **(G)** U937 cells. Human MLKL expression was induced with doxycycline (Dox) and treated with necroptotic stimulus (TNF, Smac mimetic, IDN-6556; TSI) in the presence or absence of MLKL inhibitor necrosulfonamide (NSA; 1 µM). Cell death was measured every hour for 24 hours by percentage of SYTOX Green positive cells quantified using IncuCyte SX5 or S3 live cell imaging. Independent cell lines were assayed in *n=2-9* experiments, with errors bars indicating the mean ± SEM. **(H, I)** Western blot analyses of whole cell lysates of doxycycline induced WT or *MLKL^-/-^* HT29 **(H)** or U937 **(I)** cells expressing *MLKL^WT^*, *MLKL^S132P^* or *MLKL^TSEE^*. **(J)** Blue-Native PAGE crude cytoplasm fractions of *MLKL^-/-^* HT29 cells under TSI stimulation (0-6 hours) in the presence or absence of NSA. **(K)** Western blot analyses of whole cell lysates taken 5 h post TSI stimulation in the presence or absence of NSA from doxycycline induced *MLKL^-/-^* HT29 cells expressing *MLKL^WT^*, *MLKL^S132P^*, *MLKL^S83D^* or *MLKL^S83D,S132P^*. **(L, M)** Evaluation of necroptotic signaling in WT **(L)** or *MLKL^-/-^* **(M)** HT29 cells expressing *MLKL^WT^*, *MLKL^S132P^*, *MLKL^S83D^*, *MLKL^S83D,S132P^*, *MLKL^S83C^, MLKL^R82S^* or *MLKL^R82S,S132P^*. Human MLKL expression was induced with doxycycline (Dox) and cells treated with TSI in the presence or absence of NSA. Cell death was measured every hour for 23 hours by percentage of SYTOX-green positive cells quantified using IncuCyte SX5 or S3 live cell imaging. Independent cell lines were assayed in *n=4* experiments, with errors bars indicating the mean ± SEM. **(N)** Western blot analyses of whole cell lysates taken in the presence or absence of 3 h post TSI stimulation from doxycycline induced *MLKL^-/-^* HT29 cells expressing *MLKL^WT^*, *MLKL^S132P^*, *MLKL^S83C^, MLKL^R82S^,* or *MLKL^R82S,S132P^.* Blots images in **B, H, I, J, K** & **N** are representative images of at least two independent repeat experiments.

**Supplementary Figure 2.**
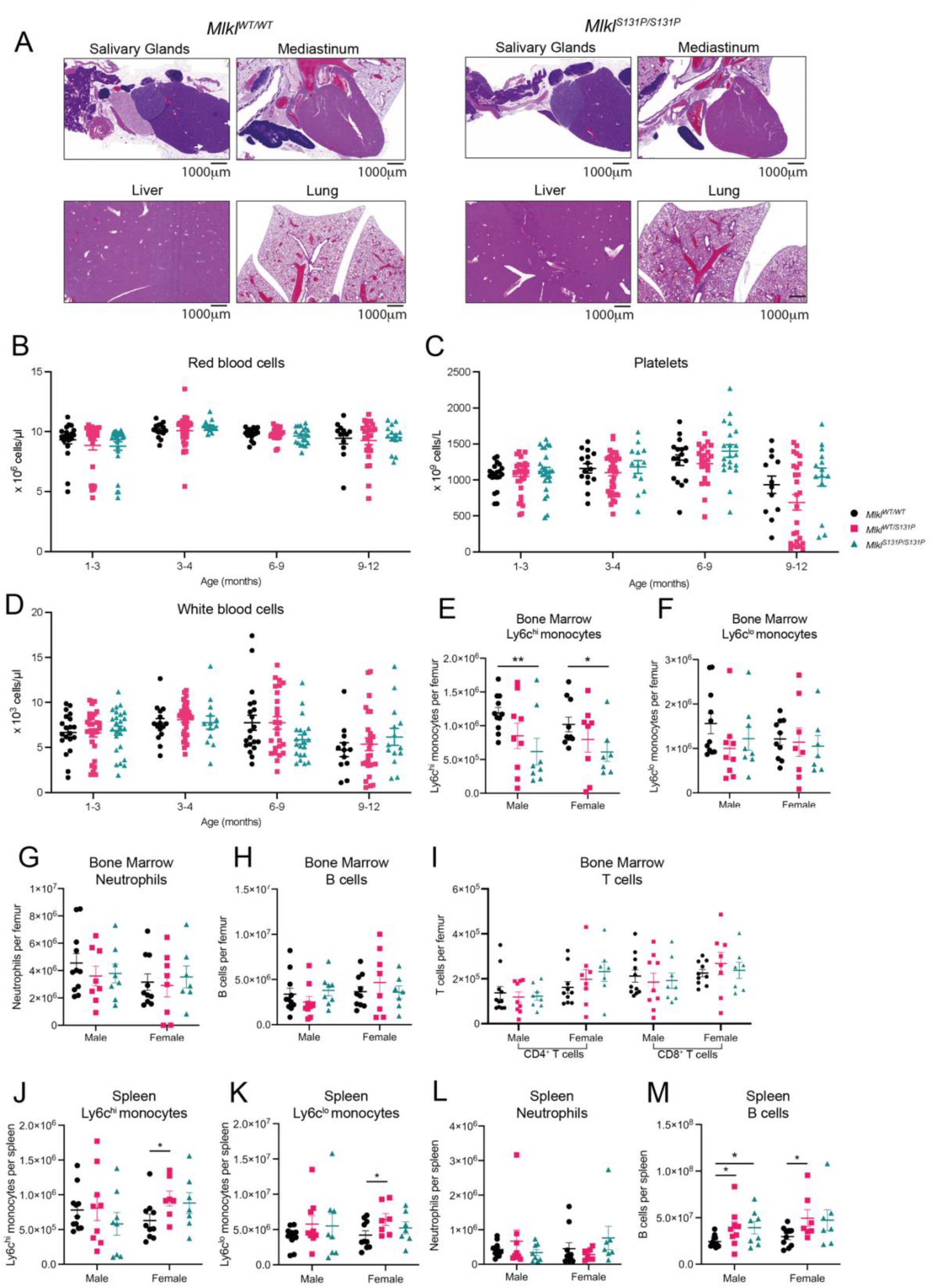
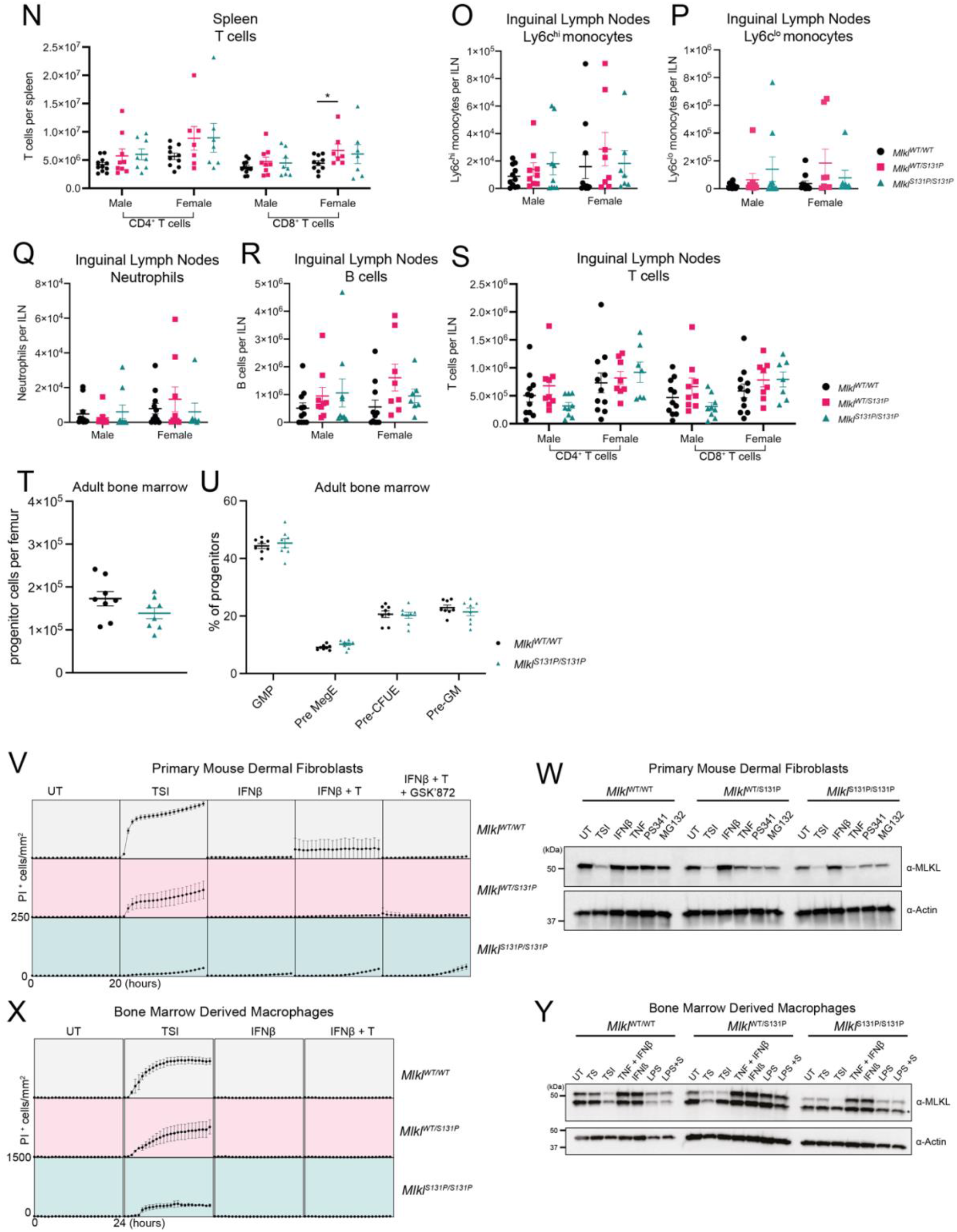
Cells expressing endogenous MLKL^S131P^ exhibit reduced TSI-induced necroptotic cell death. **(A)** Representative images of H&E staining of salivary glands, mediastinum, liver and lung from 7–9-month-old *Mlkl^WT/WT^* and *Mlkl^S131P/S131P^* mice. Images are representative of *n=2* mice per genotype. **(B-D)** ADVIA hematology quantification of circulating red blood cells **(B)**, platelets **(C)** and white blood cells **(D)** in *Mlkl^WT/WT^*, *Mlkl^WT/S131P^* and *Mlkl^S131P/S131P^* mice across age. Each symbol represents one independent mouse sampled and error bars represent mean ± SEM for *n=12-38* mice as indicated. (**E-S)** Flow cytometry quantification of innate (Ly6C^hi^, Ly6C^lo^ and neutrophils) and adaptative (B cells, CD4^+^ T cells and CD8^+^ T cells) immune cells in the bone marrow **(E-I)**, spleen **(J-N)** and inguinal lymph nodes **(O-S)** of 8–12-week-old basal state *Mlkl^WT/WT^, Mlkl^WT/S131P^,* and *Mlkl^S131P/S131P^* mice as indicated. Each symbol represents one independent mouse sampled and error bars represent mean ± SEM for *n=7-11* mice as indicated. **(T, U)** GMP, MegE, CFU-E, and Pre-GM progenitor populations in adult bone marrow were gated according to previously published strategies and presented as percentage of gated progenitors (Lin^-^cKit^+^Sca1^-^). Data presented mean ± SEM of *n* = 8, with each symbol representing an individual mouse sampled. **(V, W)** Primary mouse dermal fibroblasts (MDFs) were isolated from *Mlkl^WT/WT^, Mlkl^WT/S131P^,* and *Mlkl^S131P/S131P^*mice and stimulated as indicated for 6 hours for western blot analysis **(V)** or 20 hours for quantification of PI-positive cells using IncuCyte S3 live cell imaging **(W)**. **(X, Y)** Bone marrow derived macrophages were isolated from *Mlkl^WT/WT^, Mlkl^WT/S131P^,* and *Mlkl^S131P/S131P^* mice and stimulated on day 6 of culture as indicated for 6 hours for western blot analysis **(X)** or 24 hours for quantification of PI-positive cells using IncuCyte S3 live cell imaging **(Y)**. *p<0.05 **p<0.01 calculated using an unpaired, two-tailed Students t-test.

**Supplementary Figure 3.**
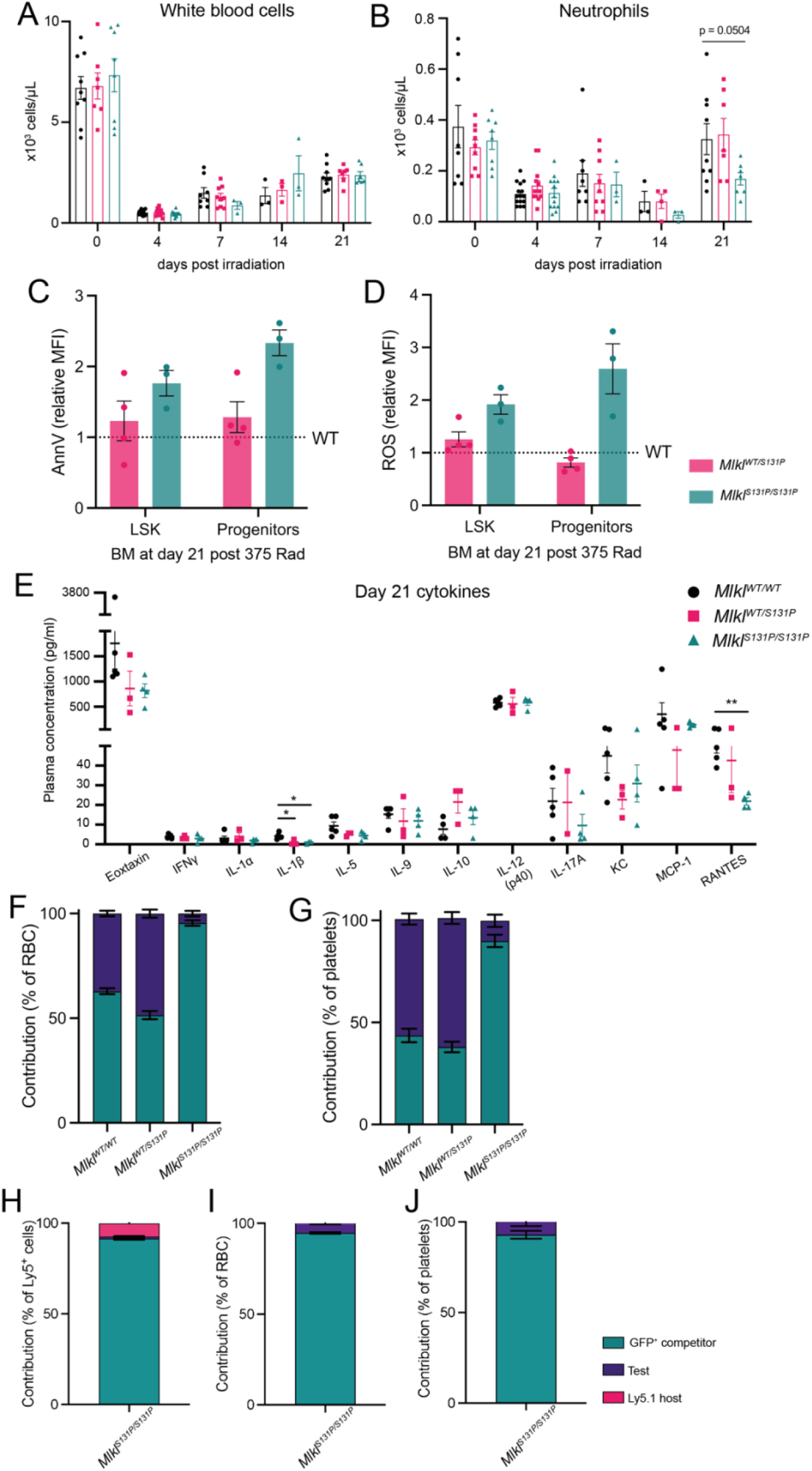
*Mlkl^S131P/S131P^* hematopoietic stem cells transplanted in excess are outcompeted at 6-weeks post-transplant. White blood cells **(A)** and neutrophils **(B)** in *Mlkl^WT/WT^, Mlkl^WT/S131P^,* and *Mlkl^S131P/S131P^* mice following treatment with 5.5 Gy radiation. Mean ± SEM of *n=*3-15 independent mice from three separate experiments. Relative amount of Annexin V **(C)** and ROS **(D)** in *Mlkl^WT/S131P^* and *Mlkl^S131P/S131P^* LSK and progenitor cells was determined 21 days post irradiation. MFI calculated relative to mean of n= 6 *Mlkl^WT/WT^* LSK and progenitor cells. Mean ± SEM of *n=*3-4. **(E)** Multiplex measurement of plasma cytokine levels at 21-days post myelosuppressive radiation. Mean ± SEM of *n=*2-4. Bone marrow from *Mlkl^S131P/S131P^* mice on CD45^Ly5.2^ background was mixed with wild-type GFP^+^ competitor bone marrow on a CD45^Ly5.2^ background at a 50:50 **(F, G)** or 70:30 **(H-J)** ratio and transplanted into irradiated CD45^Ly5.1^ recipients. Relative donor contribution to red blood cells **(F, I)**, platelets **(G, J)** and PBMCs **(H)** was assessed at 6 weeks post-transplantation. Mean ± SEM shown of n=5-11, with each donor bone marrow placed into 2-3 recipients. Host contribution (CD45^Ly5.1^) depicted in pink, GFP competitor in green, and test (*Mlkl^WT/WT^, Mlkl^WT/S131P^, Mlkl^S131P/S131P^*) in purple. *p<0.05 **p<0.01 calculated using an unpaired, two-tailed Students t-test.

**Supplementary Figure 4.**
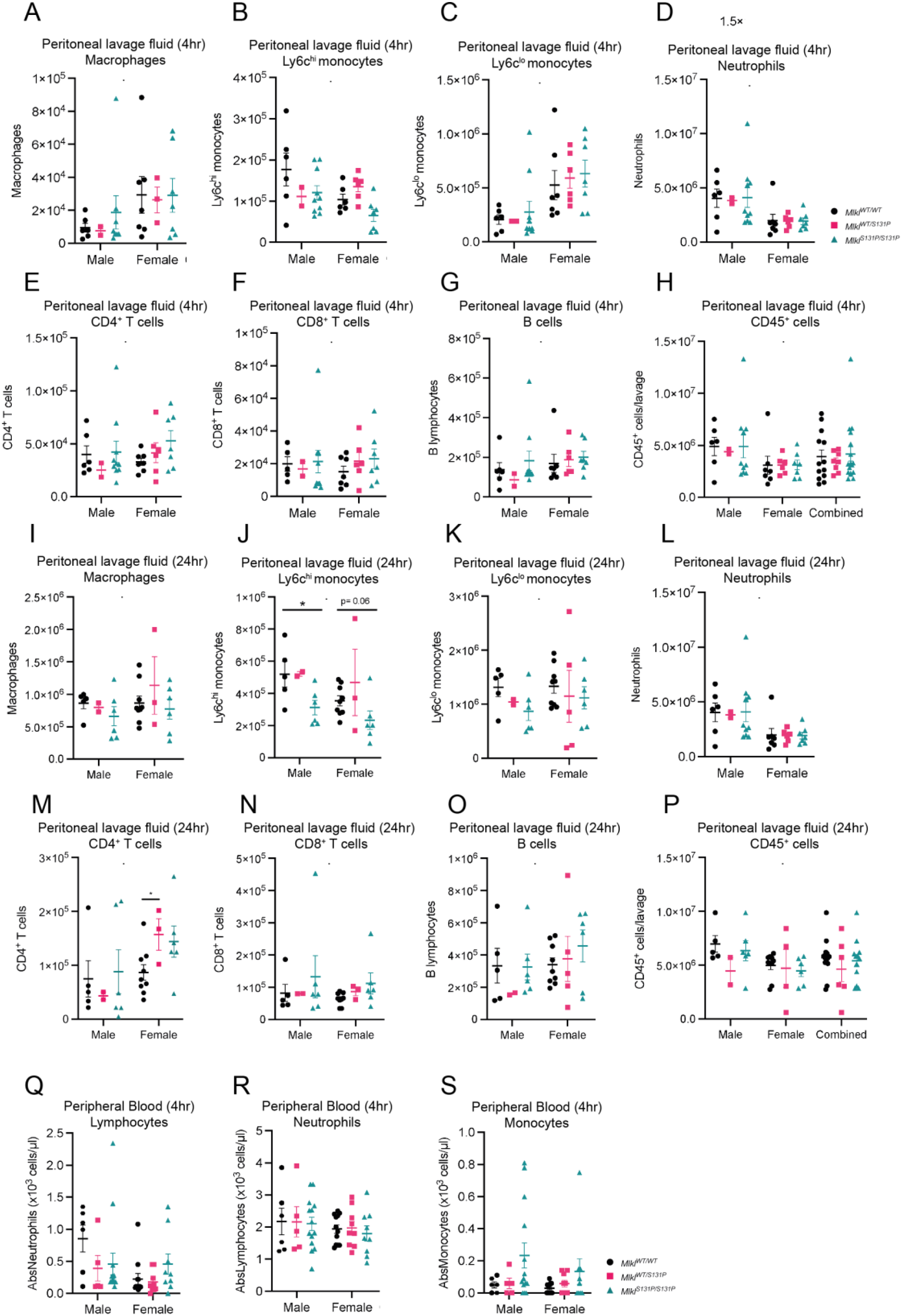
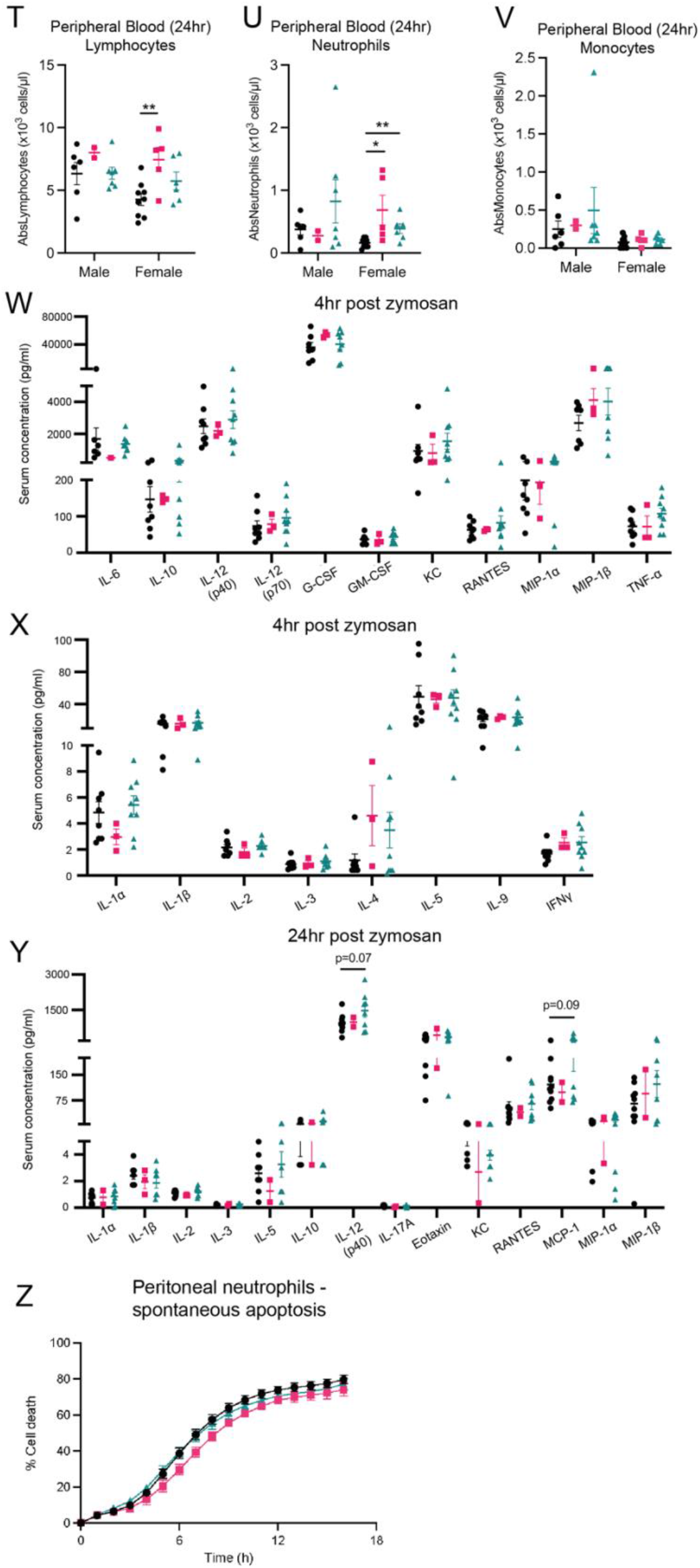
*Mlkl^S131P^* mice have reduced levels of Ly6C^hi^ monocytes in the peritoneal lavage at 24-hours post zymosan injection. **(A-P)** Flow cytometry quantification of peritoneal innate (macrophages, Ly6C^hi^, Ly6C^lo^ and neutrophils) and adaptive (CD4^+^ T cells, CD8^+^ T cells and B cells) immune cells at 4- **(A-H)** or 24- **(I-P)** hours post-intraperitoneal injection of zymosan as indicated. **(Q-V)** ADVIA hematology quantification of peripheral blood cells (lymphocytes, neutrophils, and monocytes) at 4- **(Q-S)** and 24- **(T-V)** post-injection of zymosan. **(W-Y)** Multiplex measurement of cytokine levels in peritoneal lavage at 4 **(W, X)** or 24 **(Y)** hours post-zymosan injection. Each symbol represents one independent animal, with mice from the 4- or 24-hour timepoint pooled from 3 and 2 independent zymosan experiments respectively. Error bars represent mean ± SEM for *n=2-14* mice as indicated. **(Z)** Evaluation of spontaneous apoptosis in neutrophils recruited and isolated from the peritoneum 4 hours post-zymosan injection. Neutrophils were left unstimulated (spontaneous apoptosis) for 16 hours and cell death was measured every hour by percentage of SYTOX Green positive cells quantified using IncuCyte SX5 live cell imaging. Data were collected from one independent experiment with male and female data pooled, neutrophils isolated from independent mice with mean ± SEM of *n=*6-14 presented. *p<0.05 **p<0.01, ***p<0.001 calculated using an unpaired, two-tailed Students t-test.

**Supplementary Figure 5.**
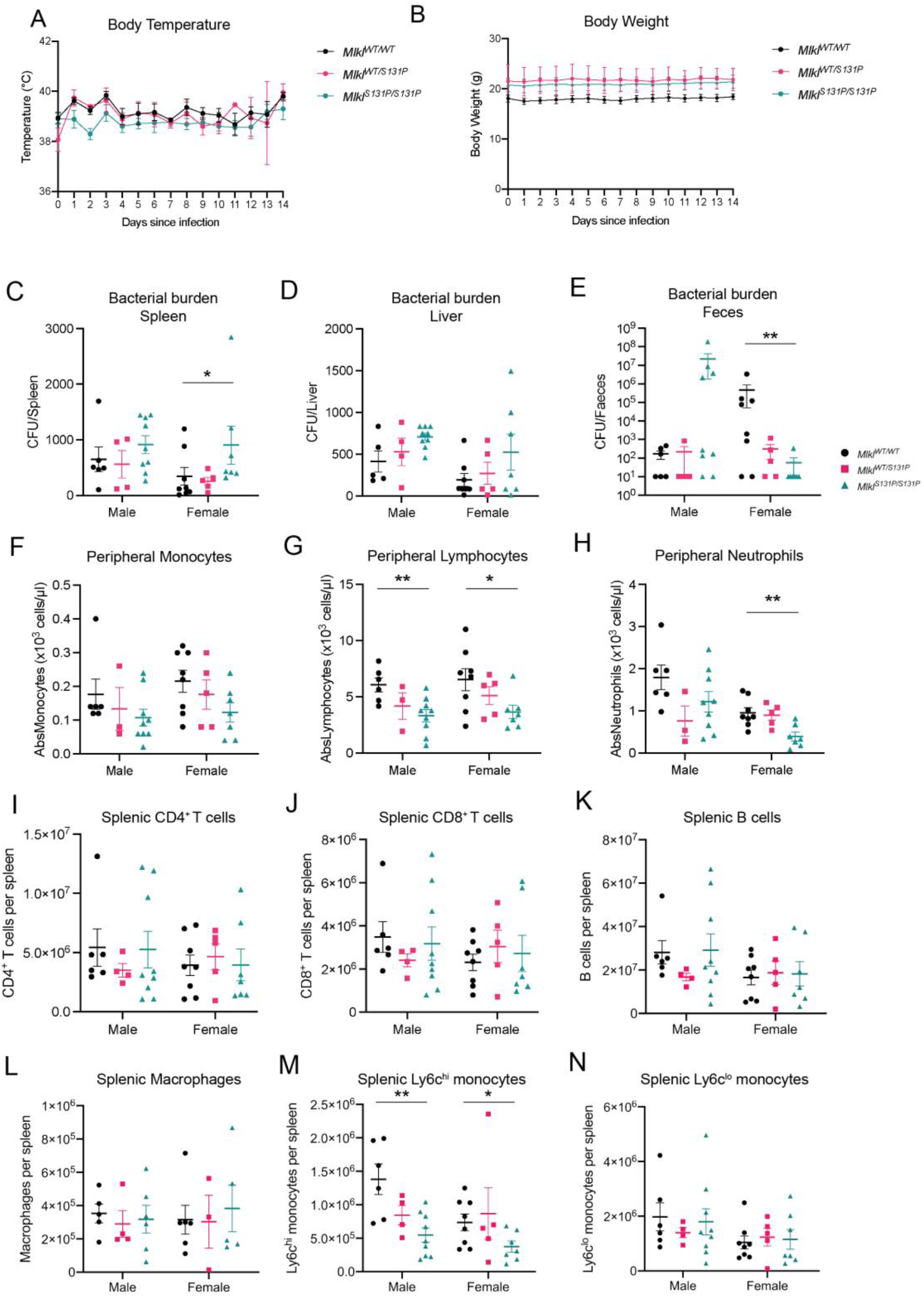
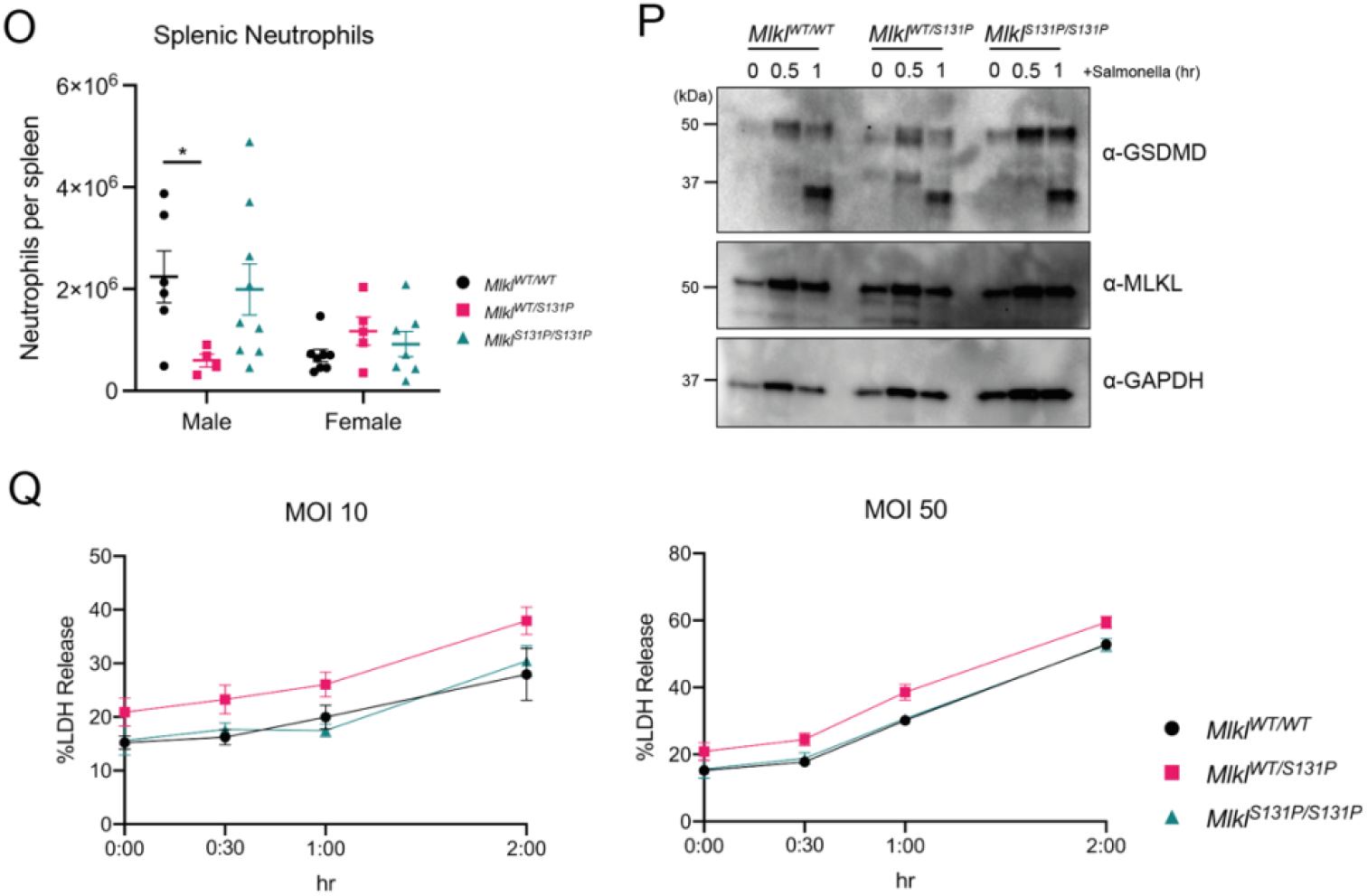
*Mlkl^S131P^* does not abrogate death of BMDMs upon *in vitro Salmonella* infection. **(A, B)** *Mlkl^WT/WT^, Mlkl^WT/S131P^,* and *Mlkl^S131P/S131P^*mice were infected with *Salmonella* via oral gavage and monitored for 14-days by daily body weight **(A)** and temperature **(B)** measurements. (**C-E)** Bacterial burden calculation of *Salmonella* colonization in the spleen **(C)**, liver **(D)**, and feces **(E)** at experimental endpoint. **(F-H)** ADVIA hematology quantification of peripheral monocytes **(F)**, lymphocytes **(G)** and neutrophils **(H). (I-O)** Splenic adaptive (CD4^+^ T cells, CD8^+^ T cells and B cells) **(I-K)** and innate (macrophages, Ly6C^hi^, Ly6C^lo^ and neutrophils) **(L-O)** immune cells were quantified by flow cytometry at experimental endpoint. *Salmonella* infection was completed 3 independent times, with each symbol representing an individual mouse sampled. Error bars represent mean ± SEM for *n=3-9* mice as indicated. **(P, Q)** *In vitro* assessment of *Salmonella* SL1344 infection of primary BMDMs generated from *Mlkl^WT/WT^, Mlkl^WT/S131P^,* and *Mlkl^S131P/S131P^* mice. **(G)** BMDMs were infected with *Salmonella* (MOI:25) and cleavage associated with Gasdermin-D activation during pyroptosis was analyzed by immunoblotting at the indicated time points. **(H)** LDH release cell death assay of BMDMs after infection with *Salmonella* (MOI:10 or MOI:50) at indicated time points. *In vitro Salmonella* experiment completed once, with mean ± SEM for *n=3* individual mice shown. Blot images in **(G)** are representative of independent duplicates.

